# A biomechanical switch regulates the transition towards homeostasis in esophageal epithelium

**DOI:** 10.1101/2021.02.03.428820

**Authors:** Jamie McGinn, Adrien Hallou, Seungmin Han, Kata Krizic, Svetlana Ulyanchenko, Ramiro Iglesias-Bartolome, Frances J. England, Christophe Verstreken, Kevin J. Chalut, Kim B. Jensen, Benjamin D. Simons, Maria P. Alcolea

## Abstract

Epithelial cells are highly dynamic and can rapidly adapt their behavior in response to tissue perturbations and increasing tissue demands. However, the processes that finely control these responses and, particularly, the mechanisms that ensure the correct switch to and from normal tissue homeostasis are largely unknown. Here we explore changes in cell behavior happening at the interface between postnatal development and homeostasis in the epithelium of the mouse esophagus, as a physiological model exemplifying a rapid but controlled tissue growth transition. Single cell RNA sequencing and histological analysis of the mouse esophagus reveal significant mechanical changes in the epithelium upon tissue maturation. Organ stretching experiments further indicate that tissue strain caused by the differential growth of the mouse esophagus relative to the entire body promotes the emergence of a defined committed population in the progenitor compartment as homeostasis is established. Our results point to a simple mechanism whereby the mechanical changes experienced at the whole tissue level are integrated with those “sensed” at the cellular level to control epithelial cell behavior and tissue maintenance.

## Main

Epithelial tissues have long been thought to have a defined catalogue of cell fate choices that can be made upon division. However, recent reports have challenged this traditional concept by revealing that cell state is not always restricted. Instead, cells possess a remarkable ability to adapt their programme of cell fate in response to varied tissue demands, such as developmental cues and damage^1–7^. This adaptability represents a clear evolutionary advantage to ensure fast and efficient cellular responses to tissue challenges. Yet, if dysregulated, this promiscuous behavior can promote epithelial disease and cancer^8^.

The mouse esophagus is an ideal model to study epithelial cell dynamics at the whole tissue level due to its simple architecture, defined size, as well as its amenity for whole-tissue 3D imaging^9^. The esophagus is lined by a squamous keratinized epithelium, formed by layers of keratinocytes (**Fig. 1a**). Proliferation is restricted to progenitor cells confined within the basal layer. Upon commitment to differentiation, basal cells exit the cell cycle and stratify into the suprabasal layers, migrating to the tissue surface from which they are shed. Using quantitative cell fate analysis in mice, it has been proposed that the esophageal epithelium (EE) is maintained by a single, functionally equivalent, progenitor population that balances the production of proliferating and differentiating cells through stochastic symmetric and asymmetric divisions^10^.

**Fig. 1:**
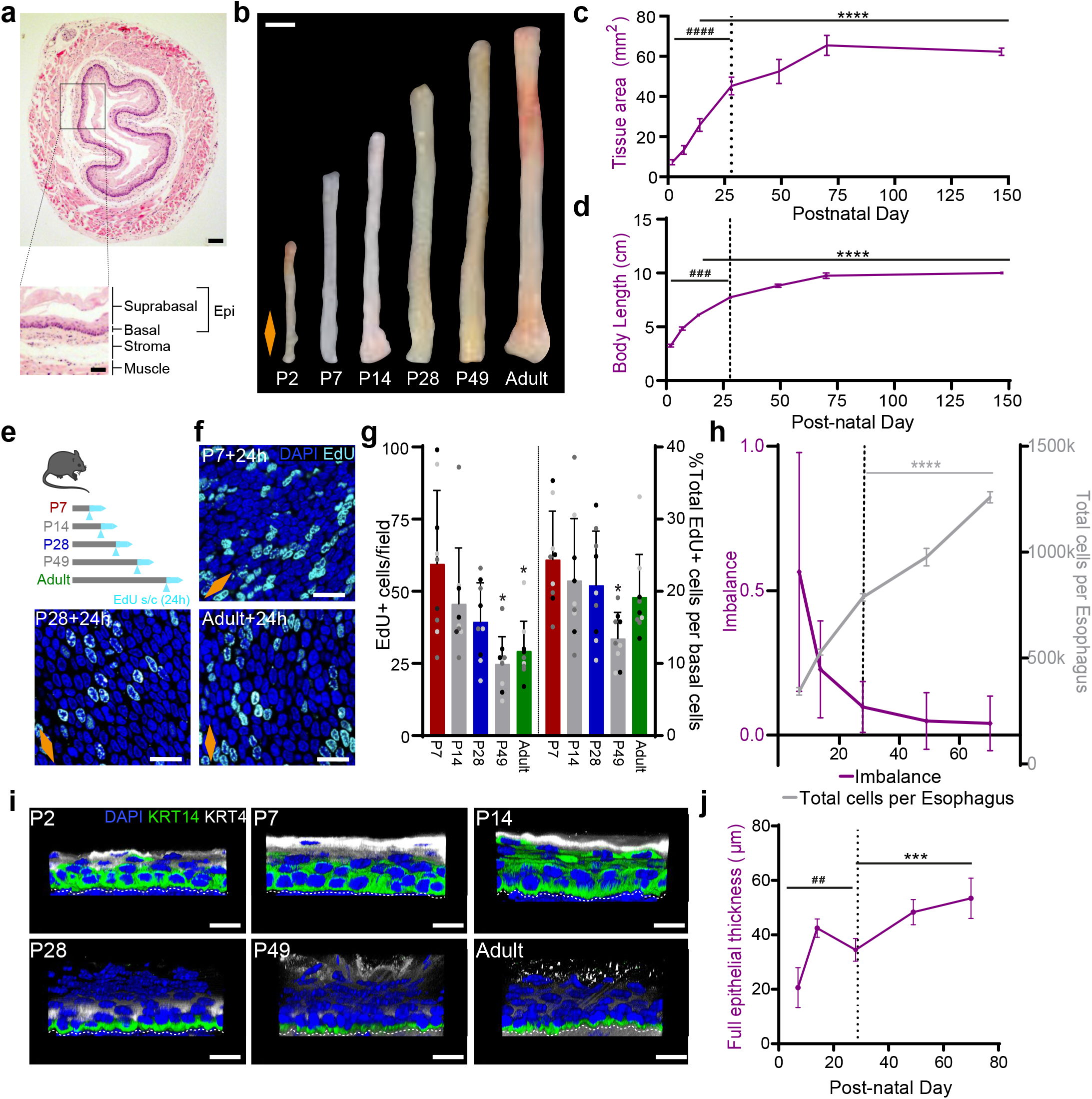
The postnatal mouse esophagus as a model of rapid tissue expansion transitioning towards homeostasis. **a,** H&E section of adult esophagus. Inset showing tissue layers. Epi, epithelium. **b,** Images of whole esophagus throughout postnatal development. **c,** Tissue area after longitudinally opening the esophageal tube. n=103; Millimeters, mm. **d,** Whole body growth in length excluding tail. n=26; Centimeter, cm. **e,** *In vivo* protocol. Mice were treated with a single EdU injection 24h prior tissue collection at the time points indicated. **f,** Basal confocal view of EdU+ cells in typical EE wholemounts from **(e)**. **g,** Quantification of EdU+ basal cells per field (left), and relative to the number of DAPI+ basal cells (right) from **(e)**. Presented as mean ± SD. Individual points show individual measurements, greyscale indicates values from each of 3 mice. n = 3. **h,** Fraction of cell fate imbalance, showing the degree to which basal cell fate is biased towards duplication over differentiation and loss (left axis; see Methods). Total cell production throughout postnatal development (right axis; see Methods). n=3. **i,** Representative 3D rendered confocal z-stacks showing side views of EE wholemounts throughout postnatal development. Dashed white lines indicate basement membrane. **j,** Epithelial thickness, including enucleated terminally differentiated layers. n=3; Micrometer, μm. All data derived from wild-type *C57BL/6J* mice, expressed as mean values ± SEM, unless otherwise stated. Data analysis was performed using one-way ANOVA with Tukey’s multiple comparisons test (##p < 0.01, ###p < 0.001 ####p < 0.0001 relative to P70; *p < 0.05; ***p < 0.001, ****p < 0.0001 relative to P7). **Scale bars**. 1A(200 μm), inset(100 μm); 1B(2mm); 1F,I(20μm). **Stainings**. Blue, 4’,6-diamidino-2-phenylindole (DAPI); cyan, EdU; green, basal marker KRT14; greyscale, differentiation marker KRT4. Dashed line in graphs indicates P28. Orange diamonds depict the longitudinal orientation of the esophagus where indicated (outlined in **Extended Data Fig. 1a**). See also **Extended Data Fig. 1**.

To identify the mechanisms that underlie the switch in cell behavior, we focused on one of the first, and most important, transitions that a specified tissue faces; the establishment of homeostasis. In many tissues, development continues after birth with an extremely rapid but physiological level of expansion that encompasses the postnatal growth of the animal, and culminates in the establishment of homeostasis^11, 12^. Once the homeostatic steady state is achieved, adult epithelial cells have been shown to retain the ability to repurpose certain developmental features as a mechanism to promote their plasticity and enhance their regenerative capacity^6, 13, 14^. These observations reveal the crucial role of developmental pathways beyond their typical developmental setting, contextualizing adult cell behavior. Yet, despite their critical relevance, the mechanisms governing the cellular responses underlying developmental transitions to and from homeostasis remain unclear.

Over the last decade, increasing evidence has emerged supporting the notion that cell behavior is not solely regulated by classically known biochemical cues. Instead, cells are able to alter their behavior in response to a combination of molecular and mechanical stimuli that translate changes taking place in the immediate neighborhood and microenvironment of a cell^15, 16,17–26^. This is exemplified during postnatal development, when individual tissues grow at their own particular pace generating significant levels of mechanical stress at the organ level^25, 27^. In this context, physical forces influence cell adhesion, shape, as well as niche availability, and, in return, feedback on tissue growth, organization and maturation^15, 28^.

Here we interrogate the postnatal development of the mouse esophagus as a physiological model to understand the mechanisms controlling changes in cell behavior at the interface between birth and adult homeostasis. Single-cell transcriptional profiling at different postnatal days provides evidence of a significant interplay between the cells of the EE and their physical environment. *In vivo* characterization shows that the developing esophagus grows at a slower pace than the rest of the body, something that results in the build-up of tensile stress along its longitudinal axis. Tissue stretching experiments further reveal that the gradual increase in physiological strain experienced by the maturing esophagus promotes the expression of KLF4 in the basal progenitor cell compartment, marking an early committed population. This mechanical transition denotes the tilt of progenitors from a proliferative expansion mode towards a balanced commitment state, a defining characteristic of adult homeostasis^10^. Our results reveal the central role that naturally occurring changes in tissue mechanics have in orchestrating the establishment of tissue homeostasis.

## Results

### The postnatal mouse esophagus as a model of rapid tissue expansion transitioning towards homeostasis

To study how epithelial cells transition in response to changes in tissue demands, we investigated the postnatal development of the esophageal epithelium (EE) and looked for changes associated with the establishment of adult homeostasis. We started by measuring changes in tissue dimensions at different time points from postnatal day (P) 2 to P147 (**Fig. 1**). To this end, esophagi from C57/Bl6 wild type (WT) mice were cut open longitudinally, and their length and width measured (**Extended Data Fig. 1b,c**). Following birth, the esophagus showed a rapid increase in surface area, reflecting the marked growth of the animal (**Fig. 1b-d and Extended Data Fig. 1b-d**). Changes in esophageal area showed a biphasic mode of growth, comprising an initial phase of rapid expansion that begins to slow down at approximately P28. By P70 tissue size stabilized (**Fig. 1b-d and Extended Data Fig. 1b-d**), coinciding with the establishment of adult homeostasis^10^.

We next looked at cell proliferation by short-term *in vivo* tracing using 5-Ethynyl-2’-deoxyuridine (EdU) incorporation assays. In this assay, a single injection of EdU labels the progenitor cells that are undergoing S phase of the cell cycle. As cells divide and complete mitosis, EdU is passed on to the daughter cells of the originally labelled progenitors marking their descendants. EdU data informs on changes in cell division rate by integrating the proportion of progenitor cells labelled in relation to their cycle time^29–31^. Quantification of the total number of EdU+ cells after a 24h chase revealed a reduction in the proliferative activity of progenitors over time, leading to a progressive decrease in the rate of basal cell production as tissue begins to reach its adult size (**Fig. 1e-h and Extended Data Fig. 1e**). The changing bias towards progenitor cell duplication is further revealed through the estimate of the degree of fate imbalance (**Fig. 1h;** see Methods). During the early postnatal period we observe a 60% imbalance towards cell duplication, which falls drastically after P14, reaching a near-balance homeostatic-like state between P28 and Adult. This data shows that the rapid basal cell proliferation found after birth stabilizes after P28.

Immunostaining for keratin 14 (KRT14), which ubiquitously marks the basal cell compartment in adult mice, was found to span all layers of the EE up until P14, demonstrating the immature profile of the early postnatal EE. The characteristic adult KRT14 profile was established by P28 (**Fig. 1i and Extended Data Fig. 1f-g**), coinciding with a switch in tissue growth (**Fig. 1c and Extended Data Fig. 1b,c**). Similarly, Keratin 4 (KRT4), typically expressed by differentiated suprabasal cells in adult EE, was found to label uneven patches of the upper-most luminal layers of the early postnatal esophagus, acquiring its typical adult distribution by P28 (**Fig. 1i and Extended Data Fig. 1g,h**). The progressive maturation of the tissue was also made apparent by an increase in the tissue thickness (**Fig. 1j**).

These results suggest that, unlike the skin, the squamous epithelium of the esophagus is not fully developed by birth^32^. Instead, our data point towards P28 as the approximate window of time when features of a matured EE become established. At this time point, we evidenced a switch away from the rapid expansion that takes place immediately after birth towards a more homeostatic mode of behavior, rendering the postnatal mouse EE as an ideal model to study the transition to homeostasis.

### KLF4 marks the emergence of an early committed population in the basal layer of the EE

Having identified the period around P28 as the window defining the switch towards homeostasis in the mouse EE, we then set out to further understand the features defining this transition. We analyzed the expression pattern of key transcription factors, known to be associated with cell fate, in immunolabelled EE wholemounts using 3D confocal imaging. These include SOX2, which has been linked to a stem/progenitor state marking the basal layer of the EE^33, 34^, and KLF4, a well characterized transcription factor marking suprabasal differentiated layers in squamous tissues^35–39^. Similar to KRT14 and KRT4 (**Fig. 1i**), Sox2 and KLF4 acquired their typical adult expression pattern by P28 (**Fig. 2a**). The uncharacteristic distribution of these transcription factors prior to P28 agrees with the postnatal EE being largely immature.

**Fig. 2:**
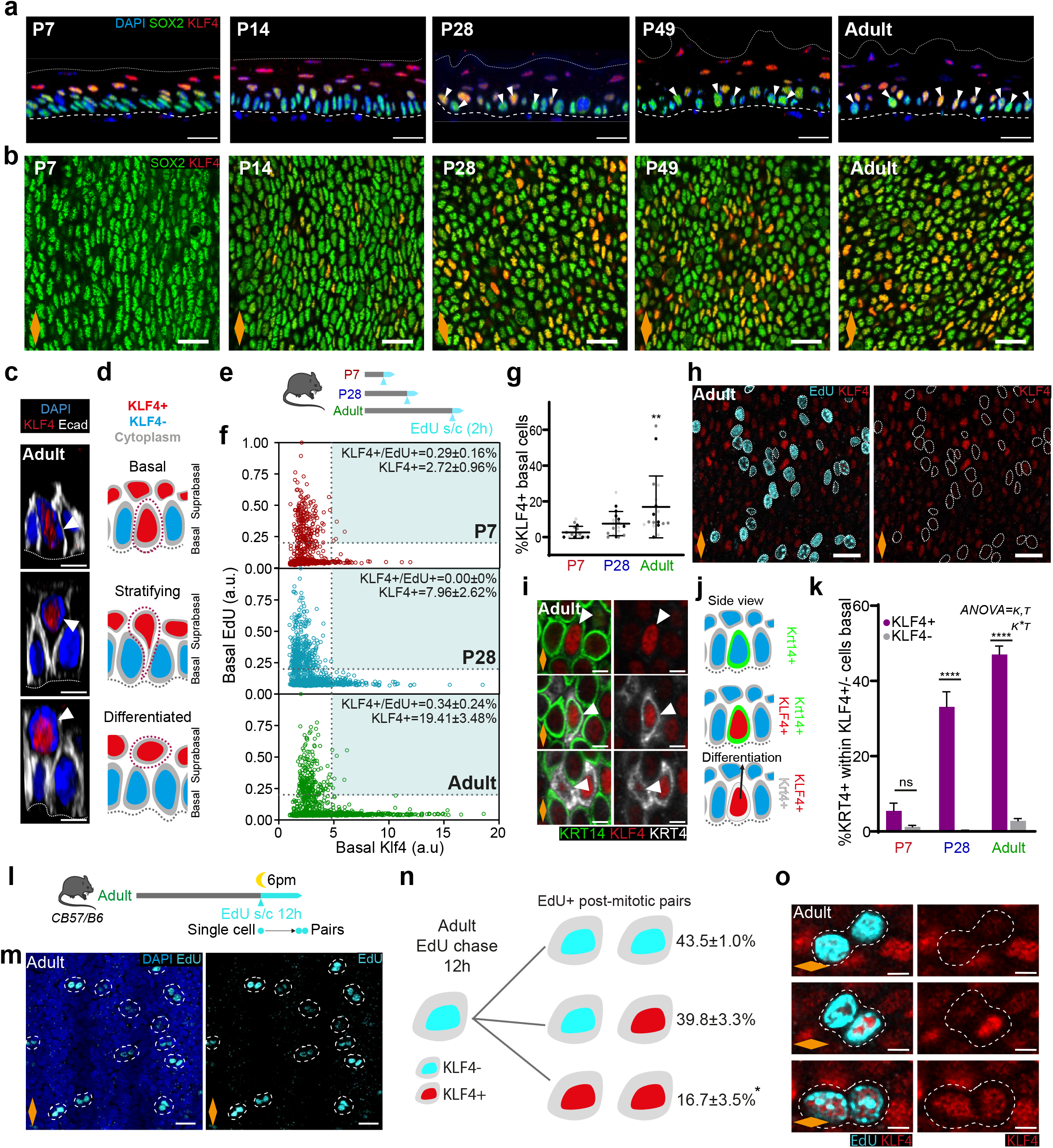
KLF4 marks an early committed progenitor population in the basal layer of the mature EE. **a,** Rendered confocal z-stacks showing side views of a typical EE wholemounts. White arrow, basal KLF4+ cells; Dashed lines, basement membrane; Dotted lines, upper limit of the EE. **b,** Representative basal views of EE wholemounts from **(a)**. **c,** Side views of 3D rendered confocal basal insets from adult EE show KLF4+ cells transitioning from basal to suprabasal layers as the cell membrane detaches from the basement membrane (dotted line). **d,** Illustration of KLF4+ (red) cells in transition from basal to suprabasal compartment as shown in **(c)**. **e,** *In vivo* protocol. Mice were treated with a single EdU injection and culled after 2 hours. **f,** Basal EdU and KLF4 intensity quantification (a.u.) from **(e)** (see Methods). n=3. **g,** Percentage of KLF4+ basal cells from **(f).** Oneway ANOVA with Tukey’s multiple comparisons test (** p < 0.01 relative to P7). Individual points show individual measurements, greyscale indicates values from each of 3 mice. n=3. **h,** Basal views of adult EE from **(e)**. Dotted lines, EdU+ basal cells. **i,** Basal views of typical adult EE wholemounts showing colocalization of KLF4 with known basal (KRT14) and suprabasal (KRT4) markers. White arrows, cell of interest. Green, KRT14; red, KLF4; greyscale, KRT4. **j,** Side view illustration depicting KLF4+ (red), KRT14 (green) and KRT4 (grey) basal colocalization combinations as shown in **(i). k,** Percentage of KRT4+ cells within KLF4+/-basal population. Data, expressed as mean values ± SEM. Two-way ANOVA with Tukey’s multiple comparisons test (****p < 0.0001; ns, not significant); n=3. **l,** *In vivo* protocol. Adult mice were administered a single dose of EdU at 6pm, and sampled after 12 hours to identify post-mitotic EdU+ basal pairs. **m,** Representative basal views of adult EE from **(l)**. Dotted white lines, post-mitotic EdU+ pairs. **n,** Schematic showing quantitative data on KLF4 expression in EdU+ pairs from **(l)**. One-way ANOVA, with Tukey’s multiple comparisons test (*p < 0.05); n=3. **o,** Basal confocal views of typical adult EE wholemounts from **(m).** Dotted white line, representative pairs. **Scale bars**. 2A,B,H, and M(20 μm); 2C,I, and O(5 μm). **Stainings**. Unless otherwise stated. Blue, DAPI; green, progenitor marker SOX2; red, differentiation marker KLF4; cyan, EdU; Greyscale, membrane marker E-cadherin (Ecad). All data derived from wild-type *C57BL/6J* mice. Orange diamonds depict the longitudinal orientation of the esophagus where indicated (outlined in **Extended Data Fig. 1a)**. See also **Extended Data Fig. 2**.

Interestingly, we noted an unreported feature of the KLF4 expression pattern in the EE. Comprehensive analysis of 3D wholemount images revealed a small, but increasing, percentage of basal cells expressing high levels of KLF4 (referred to as KLF4+), with ~9% KLF4+ at P28 increasing to ~19% in the adult EE (**Fig. 2a,b,f,g and Extended Data Fig. 2a**). This results in a “salt and pepper pattern” in the basal progenitor compartment, with Sox2+ cells interspersed with double positive Sox2+/KLF4+ cells (**Fig. 2a,b**). The temporal changes in the basal KLF4 expression profile, coinciding with the transition window at P28, identify KLF4 as a basal signature marking the transition towards adult homeostasis.

Since KLF4 is a transcription factor characteristic of differentiated cells, we reasoned that the basal KLF4+ population identified here could potentially represent a pool of early committed cells transitioning towards differentiation in the homeostatic EE (**Fig. 2c,d**). To explore this hypothesis, we initially investigated the proliferative capacity of this subpopulation using two independent approaches. First, we performed a 2h EdU-incorporation assay in WT mice at 3 critical time points spanning the esophageal growth transition: P7, representing the fastest growth phase; P28, a defined point marking the transition towards homeostasis; and Adult, as the fully homeostatic state. Basal KLF4+ cells were found to be virtually negative for EdU labelling across all time points (**Fig. 2e-f**). In a second approach we used R26Fucci2aR (Fucci2a) mice, which report cell cycle by expressing different fluorescent proteins depending on the cell cycle phase (mCherry, G1 cells; mVenus, S/G2/M cells). KLF4+ cells were absent from the dividing mVenus+ (S/G2/M) basal cell fraction of Fucci2a EE at all time points (**Extended Data Fig. 2b-e**). Collectively, these results indicate that the KLF4+ population identified in the basal cell layer of the adult EE is in a non-proliferative resting state, consistent with their potential committed status.

To further demonstrate whether basal KLF4+ resting cells were marking early committed cells in the adult progenitor compartment, we evaluated the expression profile of keratins 4 and 14 in the KLF4 bright and dim basal populations (KLF4+/-). Interestingly, we noted that KLF4+ basal cells could be found expressing any potential combination of these keratins; exclusively basal KRT14 (**Fig. 2i,** top panel), suprabasal KRT4 (**Fig. 2i,** bottom panel), or sporadically both (**Fig. 2i**, middle panel). This gradient pattern of keratin expression was a specific feature of KLF4+ basal cells, further indicating their transition towards differentiation (Schematic in **Fig. 2j**). This was more apparent upon quantification of KRT4 labelling in the KLF4 +/- basal populations (**Fig. 2k**). Approximately 47% of adult basal KLF4+ cells were also KRT4+, whilst representing less than 3% in the KLF4-basal population. This reveals a bias towards differentiation in the basal KLF4+ population. The observed KLF4+/KRT4+ bias was noticeable from P28 onwards, coinciding with the proposed P28 transition window. These results indicate that KLF4 marks a basal population transitioning from a progenitor to a differentiated state.

Collectively, our data strongly suggest that the basal expression of KLF4 defines early commitment towards differentiation. The increase in the KLF4+ basal population at P28, coinciding with tissue maturation, indicates that commitment to differentiation in the basal progenitor compartment is a signature of the transition towards homeostasis. The emergence of this committed population inevitably slows down the rapid tissue expansion that takes place immediately after birth, favoring the balanced proliferation/differentiation characteristic of the adult EE.

### KLF4 marks progenitor commitment promptly after cell division

We next took advantage of KLF4 expression as a basal marker of early differentiation to explore whether progenitor commitment is acquired concomitantly with cell division, a long-standing question in the epithelial stem cell field^40–44^. To this end, the expression of KLF4 was assessed in sister EdU+ pairs after a 12h chase in adult mice (**Fig. 2l,m**)^31^. KLF4 levels were reliably detected in EdU sister cells hours after their physical post-mitotic separation. Quantitative analysis showed that the majority of divided cell pairs were localized to the basal layer, and either remained KLF4- (~43%), or showed one KLF4+ cell (~40%). The remaining ~17% showed both nuclei bright for KLF4 (**Fig. 2n,o**). Very rarely, an EdU+ pair presented one sister localized in the immediate suprabasal layer (7 out of 150 pairs; ~5%), in which case the suprabasal cell was always KLF4+. Our pair analysis indicated that a marked fraction of cells acquires KLF4 expression at the cell division stage, supporting the notion that fate may be assigned at the time of division.

However, a substantial fraction of EdU+ pairs (2/5) remained negative for KLF4. Indeed, in order to fit our EdU pair data to previous quantitative lineage tracing analysis, reporting 80% of asymmetric basal divisions in the adult esophagus^10^, a proportion of these KLF4-pairs would need to make their commitment decision at a later stage. This would ensure that a subset of basal progenitor cells retain a certain degree of flexibility in their fate outcome in order to face specific tissue needs and potential challenges.

### Single-cell transcriptional profiling captures onset of homeostasis and reveals changes in tissue mechanics

To further define the molecular regulation governing the transition of the EE to homeostasis, we made use of large-scale transcriptomics at single cell resolution, focusing on three critical time points defining the postnatal transition: P7, P28 and Adult (**Fig. 1 and Fig. 2**).

Single-cell RNA sequencing (scRNAseq) was performed on the viable epithelial cell fraction (EpCam+/Cd45-) isolated from the esophagus of C57BL/6 wild-type mice (**Fig. 3a and Extended Data Fig. 3a**). Approximately 26,000 cells from 3-6 samples per time point were analyzed. The predominant expression of basal progenitor markers throughout the entire dataset (including Krt14, Itgb1 and Itga6) reflects an enrichment of basal cells under the experimental conditions used (**Extended Data Fig. 3b**). Indeed, increased expression of differentiation markers (such as Krt13, Krt4, Tgm3, Grhl3, and Sbsn) was restricted to a small subset of cells (**Extended Data Fig. 3b-d**). The UMAP distribution of single cell data revealed a noticeable separation of P7 cells, while P28 and Adult cluster closely together (**Fig. 3b**). This confirms that by P28, the EE resembles the adult homeostatic state. P28 and Adult time points were, therefore, grouped when exploring gene expression patterns, unless otherwise stated.

**Fig. 3:**
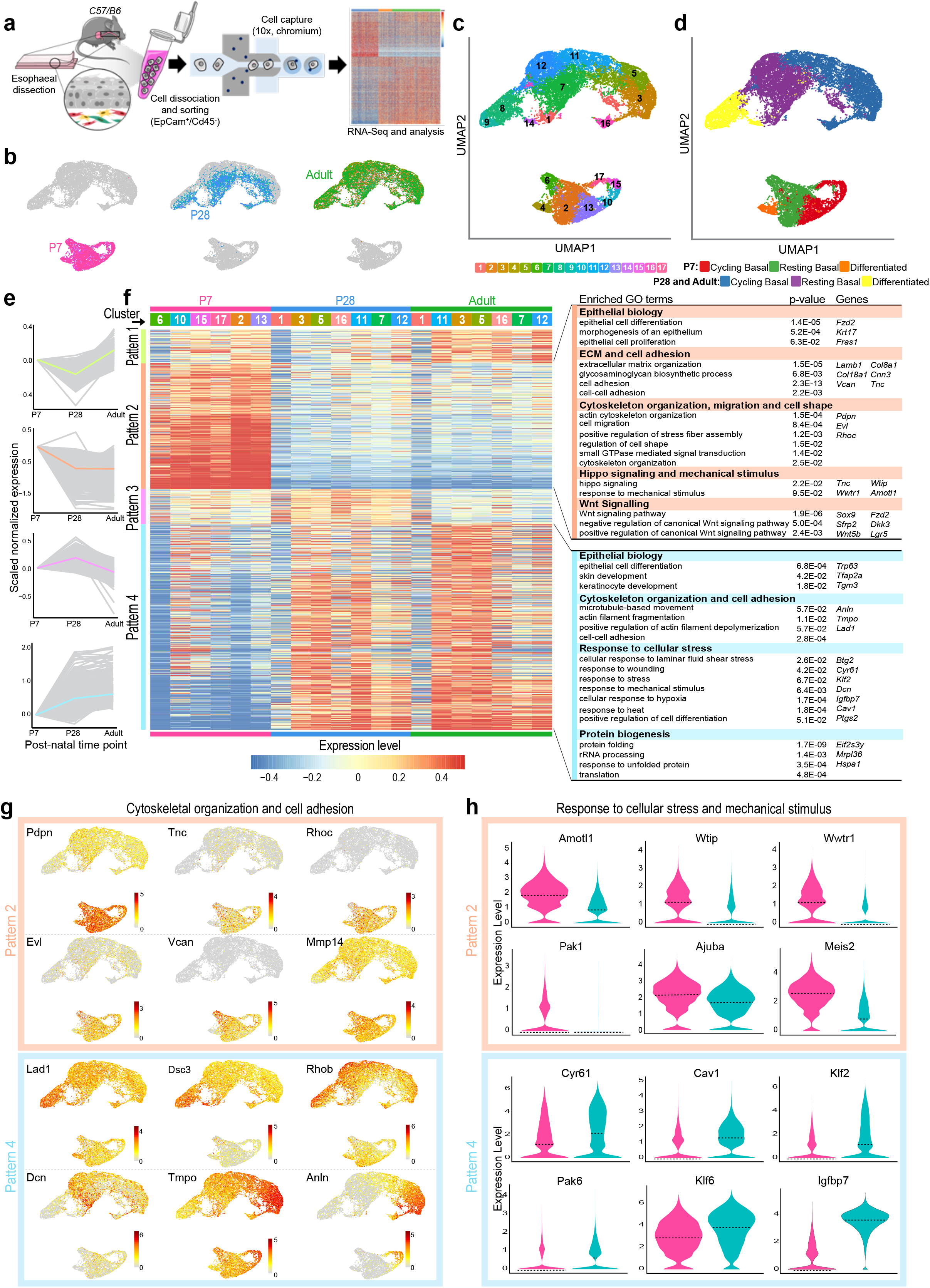
Single-cell transcriptional profiling defines transitions towards homeostasis. **a,** Schematic of single-cell RNA-seq data generation from EE sorted cells (EpCam^+^/Cd45^-^) using 10x platform. **b,** Cell distribution in the dimension reduction space UMAP at different postnatal time points. **c** and **d,** UMAPs representing cell clusters based on louvain clustering **(c)**, and annotated cell types **(d)**. **e,** Distinctive patterns (Pt) of gene expression in basal cells as defined in **(d)**. Grey, relative expression profiles of individual genes belonging to each pattern. Solid colored lines, median values at each time point. To calculate the relative expression profiles, log2-transformed normalized UMIs were scaled and averaged across all basal cells at each time point and adjusted compared to the value at P7. **f,** Heatmap representing expression of individual genes belonging to the 4 patterns in **(e)**. For expression values, log2-transformed normalised UMIs were scaled and averaged across all basal cells for each cluster and time point. The table on the right shows selected GO terms for major Pt2 and Pt4, corresponding p-values and representative genes. Closely-related GO terms are grouped together. **g,** UMAPs showing single cell expression prolife of representative genes for cytoskeletal organization and cell adhesion in Pt2 and Pt4. Color bars indicate log2-transformed normalized UMIs. **h,** Violin plots showing the expression distribution of genes related to “Response to cellular stress and mechanical stimulus” in Pt2 and Pt4. The expression level is log2-transformed normalized UMIs. All data derived from wild-type *C57BL/6J* mice.

The combined dataset was subsequently divided into 17 clusters, using an unsupervised algorithm (**Fig. 3c,** see Methods). Clusters were then annotated using signatures of known lineage and cell cycle markers (**Fig. 3d and Extended Data Fig. 3b-f**). As a result, we identified three different populations for all three time points (**Fig. 3d**): i) Actively cycling basal cell progenitors, defined by increased expression of both basal and cell cycle markers; ii) Resting/committed basal cells, with enriched basal signature expression and reduced levels of cycling markers; iii) Differentiated cells, marked by increased expression of genes associated with differentiation, as well as reduced levels of both basal and cell cycle marker genes (**Fig. 3d and Extended Data Fig. 3b-f**). Of note, although suprabasal cells at early time points do not proliferate (**Extended Data Fig. 3g,h**), they can be seen to express key markers of basal identity (**Fig. 1i and Extended Data Fig. 3b-d,f**). For this reason, clusters containing resting basal cells at P7 likely represent a combination of both basal and basal-like suprabasal cells. The basal cell populations defined in this study, including actively cycling and resting cells, are consistent with previous work on epithelial cell dynamics in the adult mouse esophagus^10^. Indeed, this study describes the basal layer as an interspersed population of cycling and early committed resting progenitors.

As expected, Klf4 expression was higher in differentiated clusters. However, in agreement with the notion that Klf4 is an early commitment marker, its expression was also enriched in one basal cluster (cluster 12) in both P28 and Adult samples (**Extended Data Fig. 4a;** middle and lower panel). Cluster 12 in the maturing EE would then represent a resting basal subpopulation enriched for Klf4 but expressing low levels of suprabasal Krt4. An equivalent cluster for this population could not be found at P7 (**Extended Data Fig. 4a;** top panel), reinforcing the idea that this early committed population is established in the basal layer as the tissue transitions towards homeostasis (**Fig. 2**). Interestingly, Klf4 mRNA expression was detected at minimal levels in all 17 clusters across samples.

Next, to shed light into the regulatory processes governing the epithelial transition towards homeostasis, we clustered genes based on their basal expression pattern over time (See Methods). Four major patterns of differentially expressed genes (DEG) at P28 and Adult relative to P7 emerged (**Fig. 3e**), with pattern (Pt) 2 and Pt4 representing the majority of DEGs. Pt2 contained genes with the strongest expression at P7 (545 genes), while Pt4 covered genes enriched towards an Adult homeostatic signature (892 genes). We performed Gene Ontology (GO) enrichment analysis for the genes belonging to each major pattern. Pt1 and 3 significantly overlapped in their GO signatures with Pt2 and Pt4, and so focus was placed on the latter. The GO analysis for Pt2 revealed particularly strong WNT signature (Wnt5b, Fzd2, Sox9, Sfrp2, Lgr5, Dkk3, and Wnt11) at P7 (**Fig. 3f and Extended Data Fig. 4c**). Other signaling genes with notable differences included increased expression of IGF related genes (Igfbp3, Igfbp4, Igf2, and Igf1r), as well as Pdgfa, Fgfr2, and Fgfr3 (**Extended Data Fig. 4c**), hinting at an active crosstalk between epithelial and mesenchymal cells at early postnatal stages^45, 46^. These transcriptional profiles reflect the increased proliferative and regenerative state observed in the esophagus shortly after birth^47–49^. Indeed, at P7, the EE showed an enrichment in genes directly associated with tissue regeneration, some of which were found to be absent in P28 and Adult (**Extended Data Fig. 4c**). Among these, we found Krt8, Krt17, Krt19 and Lgr5, associated with epithelial development, wound healing and cancer^50–52^. In contrast, genes involved in tissue maintenance and homeostasis, including Cd44, Ly6a, and Mt2^53^, were enriched at later time points (P28 and Adult), indicative of epithelial maturation. Pt4 also reflected this maturation process, drifting towards a BMP-driven regulation in P28 and Adult cells (Bmp3, Bmp4, Btg2, Hoxc4, Hoxc5, Hoxc8) (**Fig. 3f and Extended Data Fig. 4d**). This is consistent with cells favoring differentiation over expansion as the EE establishes homeostasis^54, 55^. Further Pt4 enriched GO terms indicate metabolic changes and increased protein biogenesis (**Extended Data Fig. 4d**), processes known to be associated with epithelial differentiation and maturation^56, 57^.

We identified a subset of related GO terms found to significantly impact both patterns, Pt2 and Pt4. We interpreted these shared changes as an indication of an active switch, or a transitory process. These included GO terms associated with extracellular matrix (ECM) and cytoskeleton organization, cell adhesion, and response to mechanical stimulus and stress (**Fig. 3f,g**), all critical for the mechanical stability and behavior of epithelial cells^58, 59^. In particular, specific genes involved in cell adhesion, as well as cytoskeletal and ECM organization were found to be enriched at P7 (e.g. Pdpn, Tnc, Rhoc, Evl, Vcan, Mmp14), or P28 and Adult (e.g. Lad1, Dsc3, Rhob, Dcn, Tmpo, Anln), being indicative of significant tissue remodeling taking place throughout postnatal development (**Fig. 3f,g and Extended Data Fig. 4c,d**). A particular subset of genes, encoding for the main fibrous components of the ECM, i.e. collagen, laminin and fibronectin (e.g. Col18a1, Col8a1, Col4a1, Col4a2, Lamb1, Lamc2, Fbn1), showed increased expression at P7 compared to later time points (**Extended Data Fig. 4c**). Similarly, genes encoding for ECM proteins associated with wound healing, such as Tenacin-C (Tnc), were also enriched in P7 samples (**Fig. 3f,g**). Under normal adult conditions, fibroblasts synthesize most ECM components; our transcriptional data reveals an increased contribution of early postnatal keratinocytes to ECM formation, reminiscent of active epithelial regeneration^60, 61^.

Genes involved in response to stress and mechanical stimuli were also found to be represented at both P7, as well as P28 and Adult (**Fig. 3h**). Amotl1, Wtip, and Ajuba, which have been proposed to act as negative regulators of the mechanosensor pathway Hippo, showed an increased expression at P7^62, 63^. Conversely, Wwtr1, which encodes for TAZ protein, a transcriptional regulator of the hippo pathway, was enriched at P7. P28 and Adult were marked by an increase in Cyr61 expression, a direct downstream target of YAP in mouse keratinocytes, as well as other genes modulated by YAP including Ptgs2, Igfbp7, and Klf6 (**Fig. 3f-h**)^64, 65^. Cav1, a key cellular mechanoregulator also known to positively control YAP in response to substrate stiffness, showed increased expression at P28 and Adult^64, 66, 67^. Interestingly, the expression of Klf2, a shear stress-responsive transcription factor known to be critical for heart valve development, was also significantly increased at later time points^68, 69^.

Our scRNAseq analysis indicates that the typical transcriptional signature of the adult EE is established by the transition window at P28. Analysis of this dataset highlights a clear shift in cell state through postnatal development, from being highly morphogenic at P7, to a more mature state at P28 and Adult. Moreover, changes in genes associated with cytoskeletal remodeling, adhesion, and ECM point towards changes in the mechanical properties of the tissue that, in turn, appear to impact the mechanical response of epithelial cells.

### Transition to homeostasis coincides with spatial tissue reorganization

The biomechanical signatures identified in our transcriptional profiling prompted us to look for changes in the architecture of the postnatal EE that could be linked to the switch in the mechanical response of the tissue. For this, we developed a deep learning segmentation analysis pipeline (**Extended Data Fig-5a-c**). In agreement with the strong biomechanical signature identified in the scRNAseq data, we observed considerable changes in basal cell morphology, density and spatial organization of the tissue (**Fig. 4a-c**). Specifically, at early time points (P7 and P14), when proliferative activity is at its highest (**Fig. 1e-g**), basal cells are tightly packed, as evidenced by their higher cell density relative to later time points (**Fig. 4c**). At earlier time points, basal cells exhibit increased shape anisotropy, being markedly elongated along the anterior-posterior longitudinal axis of the esophagus (**Fig. 4d and Extended Data Fig. 5d,e**). This spatial anisotropy was also observed at the tissue level, as evidenced by the bidimensional structure factor of the tissue (**Fig. 4e and Extended Data Fig. 5f,g;** see Methods). Interestingly, from P28, when the tissue transitions toward homeostasis, this anisotropic pattern became less apparent. The increased cellular crowding and compressive stress experienced by the highly proliferative EE at P7 seemed to favor the orientational alignment of cells. This was supported by higher values of the nematic order parameter, which measures the average orientational order of cells in the tissue, at early postnatal stages (P7-14; **Fig. 4f;** see Methods). Alignment of densely packed cells was also consistent with previous *in vitro* and *in vivo* work^70–72^. This long-range cell alignment decreased as the EE transitions towards homeostasis from P28 onwards (**Fig. 4e,f**).

**Fig. 4:**
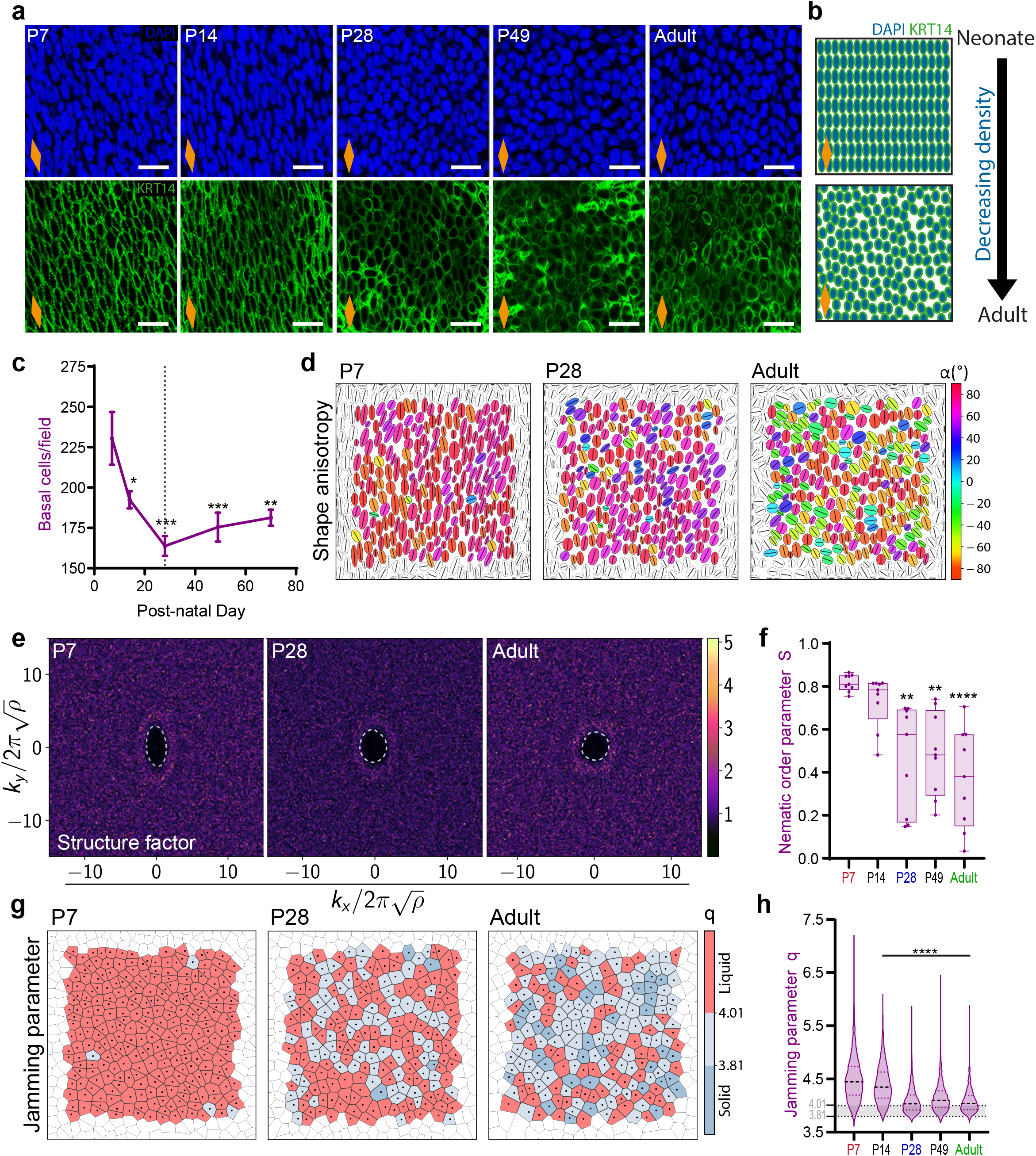
Homeostatic transition coincides with tissue spatial reorganization. **a,** Typical basal views of confocal z-stacks showing changes in cell shape, alignment and density in EE wholemounts. Blue, DAPI; green, KRT14; scale bar 20 μm. **b,** Schematic exemplifying changes seen in **(a)**. **c,** Quantification of basal cell density. Expressed as mean values ± SEM; n=3. **d,** Cell shape anisotropy tensor represented as an ellipse calculated from the nuclear centroid position of each basal cell. Long axis of each ellipse is proportional to the dominant eigen value of the tensor. Orientation is color-coded. Results from representative experiment are shown; n=3. **e,** Bidimensional structure factor quantifying basal cell spatial organization. Changes in the dashed white outline (from ellipse to circle) depict a transition from anisotropic to isotropic spatial cell distribution over time; n=3. **f,** Nematic order parameter indicative of the orientational order of cells in the tissue. Data represented as box plot; n=3. **g,** Jamming parameter (q=P/√A) represented as a Voronoi diagram calculated from the centroid of each cell. Blue, solid-like jammed-state; red, liquid-like state. Results from representative experiment are shown; n=3. **h,** Violin plot showing distribution of jammed parameter from **(g)**. Black dashed line, median; black dotted line, jamming transition value. n=2052-2594 number of segmented cells from 3 animals per time point. Voronoi diagrams are obtained from segmented 2D confocal slices of the basal layer. Nuclei at the edge of the image frame were discarded in **(e)** and **(g)** to avoid confounding effects from partially captured cells. All data derived from wild-type *C57BL/6J* mice. Data analysis was performed using one-way ANOVA with Tukey’s multiple comparisons test (*p < 0.05, **p < 0.01, ***p < 0.001, ****p < 0.0001 relative to P7; ns, not significant). Dashed line in graphs indicates P28. Orange diamonds depict the longitudinal orientation of the esophagus where indicated (outlined in **Extended Data Fig. 1a)**. See also **Extended Data Fig. 5**.

The changes observed in tissue spatial organization, cell shape, proliferation and density, are factors defining a transition from a “liquid-like” cellular organization to a non-crystalline solid-like “jammed” state. Interconversion between liquid-like and jammed states have been associated with changes in epithelial cell behavior and cell fate decisions^41, 73^. We, therefore, wondered whether this switch towards a jammed state was associated with the EE transition towards homeostasis. Using the cell shape index as a ”jamming parameter” indicator^73–75^, our data yielded two interesting observations (**Fig. 4g,h and Extended Data Fig. 5h**). First, at the whole tissue level, the basal layer of the EE appeared to transition from a liquid-like state to a more solid-like state as homeostasis is established (**Fig. 4g,h**). This is likely to be associated with the rapid cell turnover and cell crowding observed at early time points. Second, at the regional level, the adult esophagus shows a transition towards a solid-like state in the form of heterogeneous cell patches (**Fig. 4g**); a segregation proposed to result from changes in the mechanical properties of epithelial cells^76^. This data agrees with previous work, supporting the notion that jamming transitions coordinate tissue morphogenesis by coupling the global mechanical properties of the tissue with local events at the cellular scale^41, 73^.

Our results reveal how changes in the physical parameters that define epithelial tissue organization, including the long-range liquid crystalline-order, and liquid to solid-like state jamming transitions, may be directly correlated with the physiological process leading the switch from postnatal expansion to homeostasis in the EE. Thus, we propose that the transition in cell behavior towards homeostasis is guided by dynamic changes in tissue mechanics sensed at the cellular level.

### Strain generated by differential growth activates EE mechanosensing

Analysis of scRNAseq signatures together with image based characterization of tissue geometry and spatial organization suggest that changes in tissue mechanics occur during the postnatal transition towards homeostasis. We speculated that a potential source of mechanical stress may originate from growth anisotropy between the EE, neighboring tissues and/or the whole body of the animal.

To explore this scenario, we performed an immediate *in situ* post-mortem measurement of the esophageal length at different postnatal time points from P7 up to adulthood (**Fig. 5a**). The esophagus was subsequently dissected out and its *ex vivo* length measured (**Fig. 5b and Extended Data Fig. 6a**). The *in situ* and *ex vivo* lengths were compared to the mouse body length, all expressed as a percentage relative to their value at P7 (**Fig. 5c**). Naturally, the length of the *in situ* esophagus and entire body showed an almost perfect overlap. Intriguingly, the *ex vivo* length, which reflects the growth of the organ when detached from any adjacent tissues, did not follow the same trend. When approaching P28, the length of the *ex vivo* esophagus increased at a seemingly slower pace than the rest of the body, resulting in a 30% strain by adulthood (**Fig. 5d**). We confirmed that the observed discrepancy in length was not an artifact imposed by the underlying muscle layer by measuring the length of the muscle compartment independently. We found that the muscle layer was approximately 10% longer than the EE in adult samples (**Extended Data Fig. 6b,c**). Therefore, the differential growth of the developing esophagus results in the build-up of a progressive strain, estimated to be of approximately 40% in normal physiological adult conditions. We conclude that the EE undergoes a mechanical transition, from compressive stress at earlier time points, when rapid proliferation promotes cell crowding (**Fig. 4c**), to tissue stretching upon homeostasis (**Fig. 5d**).

**Fig. 5:**
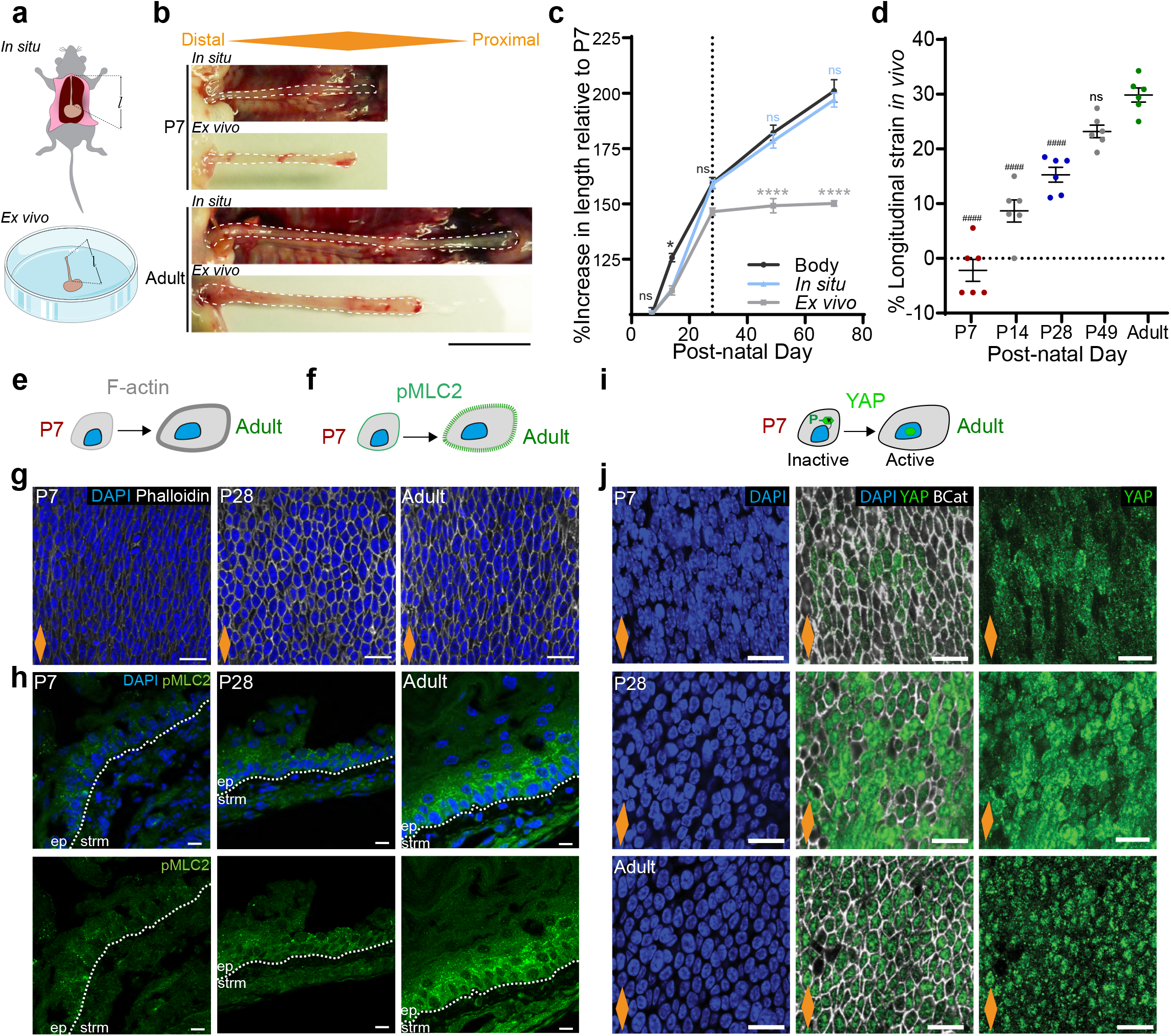
Differential growth generates longitudinal tissue strain sensed at the cellular level. **a,** Schematic exemplifying method for esophageal length measurements *in situ* and *ex vivo* (*ɭ*). **b,** *In situ* and immediate *ex vivo* images of esophageal tubes captured early in postnatal development (P7) and adult mice. White dashed lines delineate esophageal tube. **c,** Percentage increase in esophageal length (*in situ* and *ex vivo*) compared to body length (excluding tail) throughout postnatal development (P7 set to 100). Dashed line indicates P28. Data expressed as mean ± SEM. One-way ANOVA with Tukey’s multiple comparisons test (*p < 0.05, ****p < 0.0001; ns, not significant; relative to body length); n=6. **d,** Longitudinal tissue strain *in vivo* represented as percentage. Data expressed as mean ± SEM. One-way ANOVA with Tukey’s multiple comparisons test (####p < 0.0001; ns, not significant; relative to P70); n=6. **e** and **f,** Schematic representing changes in F-actin levels (grey) and pMLC2 (green) in EE cells through postnatal development. **g,** Basal views of typical EE wholemount showing Phalloidin staining for F-actin at indicated time points. Blue, DAPI; greyscale, Phalloidin. **h,** Representative side views of tissue sections showing pMLC2 staining at indicated time points. Blue, DAPI; green, pMLC2. **i,** Schematic representation of changes in YAP (green) localization during postnatal development. **j,** Basal view of representative EE wholemounts showing progressive translocation of YAP to the nucleus as tissue matures. Blue, DAPI; green, YAP; greyscale, B-Catenin (BCat). **Scale bars**. 5B(1 cm); 5G,J(20 μm); 5H (10 μm). All data derived from wild-type *C57BL/6J* mice. Orange diamonds depict the longitudinal orientation of the esophagus where indicated (outlined in **Extended Data Fig. 1a)**. See also **Extended Data Fig. 6**.

The ECM constitutes an important factor for tissue mechanics, representing the scaffolding over which epithelial cells are sustained^15^. Second harmonic generation (SHG) imaging revealed an active remodeling of ECM structural components during esophageal development (**Fig. 6d**), in line with the ECM signature identified by scRNAseq (**Fig. 3g and Extended Data Fig. 4c**). Quantitative analysis showed that the ECM mesh at P7 consists of a wide range of fiber orientations (**Extended Data Fig. 6d,e;** top panels). This profile is lost by P28, when fibers become aligned transverse to the esophageal tube (**Extended Data Fig. 6d,e**), which may represent a mechanism to dissipate part of the tensile load of the tissue along its longitudinal axis.

**Fig. 6:**
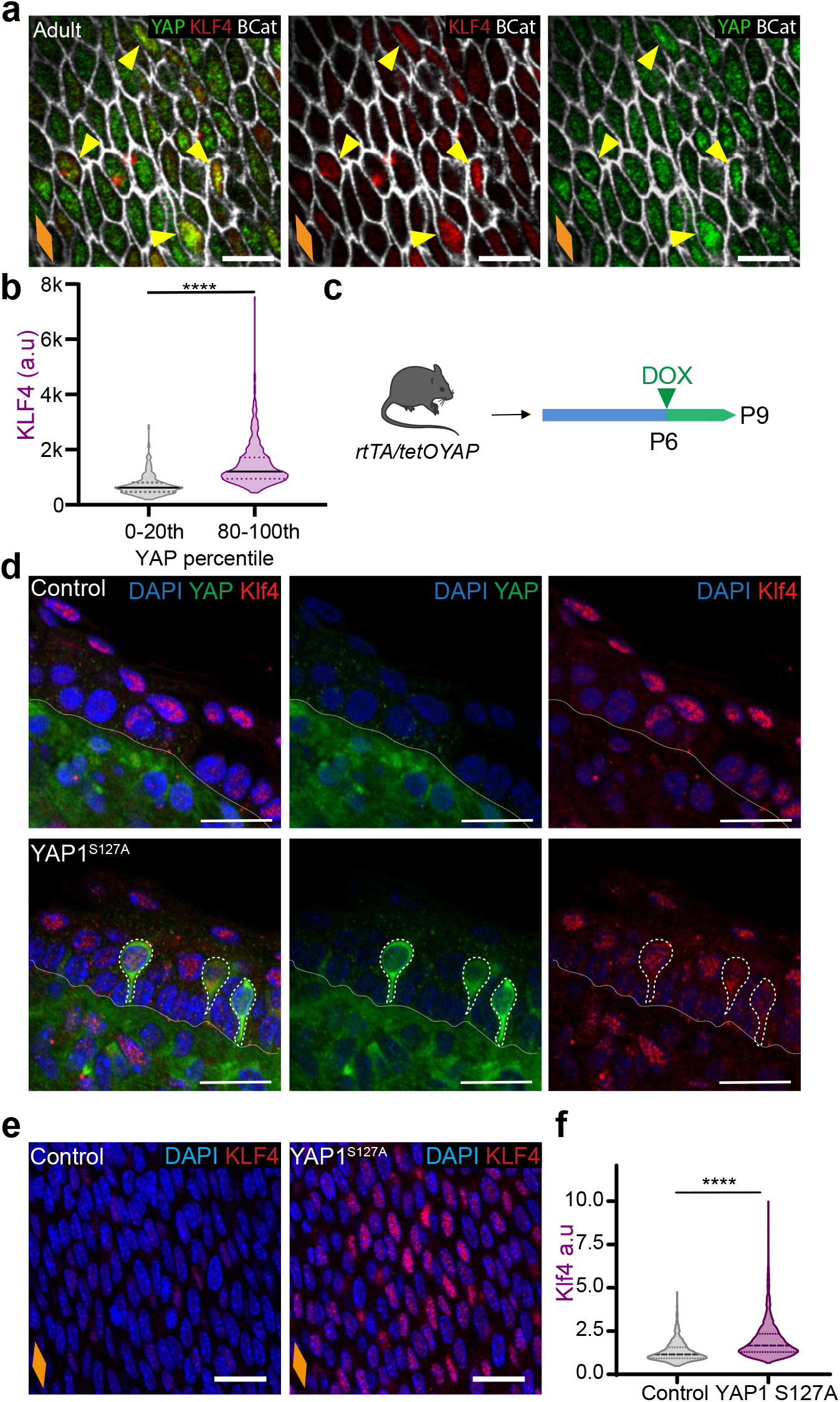
YAP nuclear localization promotes increased levels of basal commitment through KLF4. **a,** Confocal basal views of typical adult EE from *C57BL/6J* WT mice. Yellow arrows indicate colocalization of KLF4 (high) and YAP (high) staining. **b,** Quantification of KLF4 intensity (a.u) within the lowest and highest YAP expressing cells (0-20^th^ and 80-100^th^ percentile respectively). n=296 per percentile. **c,** *In vivo* protocol. P6 rtTA/tetOYAP pups were doxycycline treated for 3 days to induce an active form of YAP (S127A), and culled at P9. **d,** Staining in OCT embedded EE cryosections (10 μm thick) of YAP activated mice from **(c)**. Nuclear KLF4 expression co-localizes with YAP+ cells. Dotted white lines, YAP+ cells. **e,** Representative basal views of EE wholemounts from **(c)**. **f,** KLF4 intensity quantification (a.u) in control and YAP overexpressing mice as shown in **(e)**. n=1362-1779 cells from 3 animals. **Scale bars**. 6A(10 μm), 6D,E(20μm). **Stainings**. Blue, DAPI; green, YAP; red, KLF4; greyscale, BCat. Data were analyzed using two-tailed unpaired t test (****p < 0.0001). Orange diamonds depict the longitudinal orientation of the esophagus where indicated (outlined in **Extended Data Fig. 1a**).

To test whether the changes in the mechanical properties of the esophagus stimulated a switch in tension and cytoskeleton remodeling of epithelial cells, we labelled the EE for F actin and phosphorylated myosin light chain II (pMLC2), a marker of actomyosin contractibility (**Fig. 5e,h**). Both showed a similar profile with an increased signal from P28. We next interrogated the cytoplasmic-to-nuclear translocation of the transcription factor YAP, an effector of the mechanosensing pathway Hippo^77^. YAP nuclear localization is associated with increased space availability, cell spreading, and subsequent cytoskeletal reorganization. As predicted by the increase in tensile strain observed in the EE, YAP was found to translocate from its cytoplasmic localization at P7 to the nucleus in adult animals, with an intermediate state at P28 reflecting a mechanosensing transition (**Fig. 5i,j**). This data is indicative of basal progenitors sensing the physiological changes in strain and space availability experienced by the tissue.

We conclude that the mechanical properties of the esophagus undergo a significant change during postnatal development. This results in the physiological build-up of a longitudinal tissue tensile strain as the transition towards homeostasis takes place. The cellular response to this mechanical stimulus is evidenced by the cytoplasmic to nuclear translocation of the Hippo effector YAP.

### YAP nuclear localization promotes increased levels of basal commitment through KLF4

The experiments above suggest a temporal correlation between YAP and KLF4 expression patterns, with both transcription factors becoming localized to the nucleus of basal cells as the tissue transitions towards homeostasis (**Fig. 2 and Fig. 5i,j**). It is, therefore, tempting to speculate the existence of a potential link between YAP mechanosensing and cell commitment. To explore this possibility, we used two different approaches. First, we co-stained and quantified the levels of both proteins in the nucleus of adult basal cells (**Fig. 6a,b**). The data confirms the positive correlation between YAP and KLF4; with cells in the top 20th percentile of nuclear YAP expression having significantly higher levels of KLF4. Second, we tested the impact of YAP overexpression in the early postnatal EE, when basal cells are still negative for KLF4 (**Fig. 2a,b,g**). For this, we used the YAPS127A Doxycycline (Dox) inducible mouse strain R26-rtTA;tetO-YAPS127A (rtTA/tetOYAP). P6 rtTA/tetOYAP mice were Dox treated for 72h and the EE collected to analyze KLF4 expression in the basal layer (**Fig. 6c**). Tissue sections of the same samples indicated the presence of nuclear YAP staining in P6 induced animals as opposed to littermate non-induced controls, confirming the exogenous YAPS127A expression (**Fig. 6d**). In line with our prediction, we detected a significant increase in basal KLF4 levels upon YAPS127A induction (**Fig. 6e,f**). This experiment also denoted YAPS127A+/KLF4+ double positive cells departing from the basal compartment, and starting the journey towards differentiation (**Fig. 6d**). Collectively, co-localization and *in vivo* gain-of-function experiments strongly suggest that increased YAP nuclear localization is associated with basal cell commitment in the EE.

### Changes in tissue mechanics influence basal KLF4 expression

Having identified an increase in mechanical tensile strain throughout postnatal development, we next investigated the relevance of mechanical stress in driving the transition to homeostasis. To answer this question, we used a whole-organ *in vitro* approach utilizing a 3D printed stretching device^78^ (**Fig. 7a,b and Extended Data Fig. 7a**). This system allows for controlled uniaxial stretching by clamping the tissue at both ends, and applying a known level of strain (See Methods). Esophageal samples from P7, P28 and Adult animals were stretched along their longitudinal axis at a static strain of 40%, representing the physiological levels calculated to be present in the adult esophagus (**Fig. 5d and Extended Data Fig. 6c;** See Methods), and kept in culture for 48 hours. Tissues were then collected and the expression of KLF4 analyzed by confocal microscopy as a proxy for changes in cell commitment in the basal layer. Reassuringly, the unstretched *in vitro* controls exhibited a similar KLF4 expression profile to that observed *in vivo* (**Fig. 7c,** top panels). Moreover, adult tissues did not show differences in KLF4 expression or EdU incorporation between control and stretched samples, indicating that the experimental conditions were within the physiological range of tissue strain (**Fig. 7c,d and Extended Data Fig. 7b,c**). When P7 and P28 samples were stretched, KLF4 expression levels in the basal layer increased significantly reaching control adult values (**Fig. 7c,d**). These data suggest that the stress experienced by the adult EE promotes KLF4 expression in the basal compartment.

**Fig. 7:**
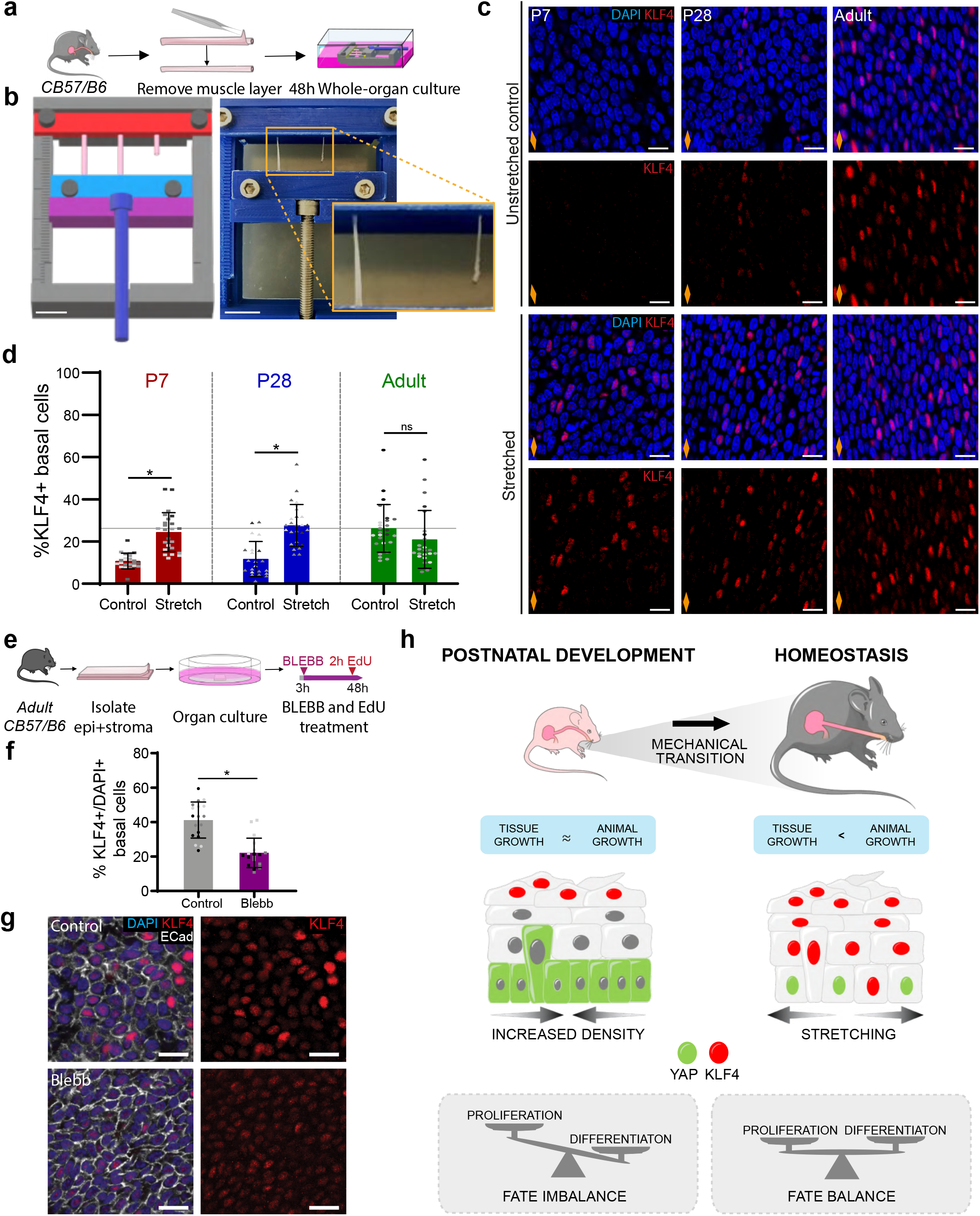
Changes in tissue mechanics influence basal KLF4 expression. **a,** *In vitro* protocol. Esophagi were collected and the muscle removed whilst maintaining tubular structure. Tissues were exposed to a 40% stretch using 3D printed stretcher and kept *in vitro* as whole-organ cultures for 48 hours. **b,** 3D model of stretching device for printing in biocompatible plastic. Left, model of stretcher. Right, device in use with adult esophagi, scale bar 1 cm. Inset on stretched and control esophagi, scale bar 5 mm. **c,** Representative basal views showing confocal images of 48 hour cultured samples in stretched and unstreched control conditions. Orange diamond indicates the longitudinal orientation of the EE and direction of stretch. **d,** Basal quantification of KLF4+ cells expressed as percentage of DAPI+ basal cells from **(c)**. n=3-6. **e,** *In vitro* protocol. Epithelial and stromal composite from adult mice were kept *in vitro* and treated with 25 μM blebbistatin (BLEBB) for 48 hours. EdU was added to media 2 hours prior tissue fixation. **f,** Percentage of KLF4+ basal cells relative to DAPI+ basal cells. n=3. **g,** Confocal basal views of typical organ cultures after a 48 hour BLEBB treatment *in vitro*. **h,** Suggested model. The differential growth of the maturing esophagus after birth results in its progressive stretching along its longitudinal axis, coinciding with a YAP nuclear re-localization. The physiological increase in strain experienced by the esophagus promotes the expression of KLF4 in the basal progenitor cell compartment, marking the emergence of an early committed population. The onset basal KLF4 expression balances proliferation, and defines the transition towards adult homeostasis. **Scale bars**. 7B(1 cm), inset (5 mm) C,G(20 μm). **Stainings**. Blue, DAPI; red, KLF4; greyscale, Ecad. All data derived from wild-type *C57BL/6J* mice, expressed as mean ± SD and were analyzed using two-tailed unpaired t test (*p < 0.05; ns, not significant Individual points show individual measurements, greyscale indicates values from each of 3 mice. Orange diamonds depict the longitudinal orientation of the esophagus where indicated (outlined in **Extended Data Fig. 1a**). See also **Extended Data Fig. 7**.

We next interfered with the tissue tension of the adult EE by disrupting actomyosin contractibility. For this, we applied blebbistatin (BLEBB), which inhibits myosin II motor activity, to esophageal organ cultures (**Fig. 7e**). *In vitro* BLEBB treatment for 48h led to an increase in cell density, without significantly impacting cell proliferation, as tested by a 2h EdU chase (**Extended Data Fig. 7d,e**). The increase in cell density is consistent with a loss of tension in the tissue. Notably, in contrast to the stretching experiment, in which we saw increased KLF4 expression, treatment with BLEBB to lower tension reduced KLF4 expression compared to untreated controls (**Fig. 7f,g**).

Taken together, our results demonstrate that the longitudinal tensile strain generated by the anisotropic growth of the esophagus during postnatal development creates a mechanical stress that tilts progenitor cells into a more committed state, balancing tissue growth and ensuring tissue maintenance upon homeostasis.

## Discussion

Here we report that the physiological strain emerging during normal postnatal development of the esophagus guides its maturation. In particular, we show that the differential growth of the esophagus relative to the entire body, from P28 onwards, generates a mechanical stretch in the tissue that influences the behavior of epithelial cells and promotes their transition towards homeostasis. This mechanical shift triggers a remarkable physical reorganization of the EE; from a densely packed, highly proliferative, basal layer to a more sparse distribution, with a decreased proliferative activity. Mechanistically, our results indicate that the naturally occurring strain experienced by the tissue after its initial expansion phase translates in YAP nuclear re-localization, and denotes a central role for KLF4 in integrating the response to these mechanical cues. The onset of basal cell commitment towards differentiation via KLF4 balances proliferation, and ultimately defines the transition to adult homeostasis.

Tissue morphogenesis has long been known to be orchestrated by finely balanced gradients of morphogens. More recent studies in a growing number of developmental systems are starting to uncover the impact that diverse types of mechanical forces have on major organs in the body^25, 28, 79^. From the shear stress dictating the development of the heart and lungs in mouse and zebrafish embryos^19, 68, 69^, to the mechanical stress guiding the patterning of the intestine and skin follicles in the developing chick and mouse^20–22, 80^. However, there is remarkably limited knowledge on the mechanisms regulating how developmental processes come to an end, safeguarding adult tissue homeostasis from abnormal growth and disease. Our findings in the postnatal mouse esophagus reveal that this transition is linked to the mechanical changes experienced by the tissue. From birth and up to P28, the esophagus expands rapidly, marked by a transient Wnt-driven regenerative transcriptional signature. After that point, the EE progressively matures and reduces its proliferative activity. However, the sustained growth of the animal at this time point inevitably results in the stretching of the esophagus, which coincides with a switch in the transcriptional signature away from Wnt signaling, and an upregulation in the expression of known stress response signals including KLFs and BMPs^68, 69, 81^. In particular, we showed that the sudden appearance of scattered basal KLF4 expression at P28 marks progenitor cells exiting from cell cycle and undergoing commitment towards differentiation, a feature defining homeostasis^10^. Indeed, our single cell transcriptional profiling identifies two coexisting basal subpopulations; actively cycling progenitors, and a resting basal population that includes KLF4 expressing cells in the adult EE. *Ex vivo* stretching experiments suggest that the mechanical stress applied to the EE is responsible for the increased KLF4 expression found in the basal cell compartment from P28 onwards. Our results demonstrate the connection between tissue strain and the acquisition of a defined committed population in the progenitor compartment upon homeostasis, and support the idea that changes in tissue mechanics can regulate epithelial behavior in response to specific tissue needs.

In agreement with previous studies, our results establish that at birth the EE presents a degree of stratification^50^. This would give the impression that cell differentiation and stratification in neonates resembles that in the adult esophagus. However, the immature features and transcriptional profile that characterize the early postnatal EE indicate that the tissue is far from homeostasis. Elevated levels of cell proliferation, increased basal cell crowding and atypical distribution of differentiation markers until P28 suggests that cell stratification in the postnatal EE may be regulated differently to the adult tissue. Indeed, a recognized feature of keratinocytes is their sensitivity to cell density, with crowding promoting a stress response leading to stratification and differentiation^31, 82–84^. Therefore, we propose that the basal cell crowding found during the initial proliferative phase in the early postnatal EE promotes a stress response that allows sufficient stratification to ensure tissue functionality up until the moment homeostasis is established.

Several *in vitro* and *in vivo* studies have shown that activation of typical mechanosensors such as YAP/TAZ and Piezo1 triggers an epithelial proliferative response^64, 85–90^. Additional work in the mouse intestinal epithelium indicates that YAP can also regulate cell differentiation, specifically through its interaction with KLF4^91^. Our findings in the postnatal EE underscore a much more complex scenario, where the mechanical stimulation of KLF4 expression may not only serve as a way to balance the production and loss of progenitor cells, but also as a mechanism to halt tissue development. These observations suggest a model in which changes in cell behavior are regulated by integrating the physiological levels of mechanical stress happening at large scale, organ-wide level, with those sensed by local neighboring cells. This mechanism provides a simple solution to orchestrate the maintenance of a whole tissue at the cellular level.

Our study offers a new perspective in understanding the regulatory processes governing epithelial cell transitions. The physiological levels of stretching, resulting from asynchronous growth of the esophagus versus the rest of the body during postnatal development, help to shape cell fate decisions and define the rules governing the maintenance of the mature tissue (See proposed model in **Fig. 7h**). This observation is likely to have a widespread impact in understanding changes in cell behavior in other mechanically active tissues, especially those where P28 also marks the point of tissue maturation, such as the alveolar lung, skeletal muscle and arteries^92–95^. Future studies should further elucidate whether regulatory processes controlled through differential growth are a broader phenomenon operating in other tissue models, as well as their relevance in regeneration and cancer^6, 96^. Indeed, the translational impact of these mechanical switches, and whether patients could benefit from therapies modulating them, represents an important area that remains to be explored.

## Methods

### Mice strains and induction of allele

All experiments were approved by the local ethical review committees at the University of Cambridge, and conducted according to Home Office project license PPL70/8866 at the Wellcome Trust – Medical Research Council Cambridge Stem Cell Institute, Cambridge University.

Unless otherwise specified, C57BL/6J mice (ordered from Charles River, UK; strain code, 632) were used for the experiments presented. Genetically engineered mouse lines used in the study include: FUCCI2a (R26^Fucci2aR^, kindly provided by Ian J. Jackson^97^), which incorporates constitutive expression of genetic probes that highlight different phases of the cell cycle (G1 marked by mCherry-hCdt1, and S/G2/M marked by mVenus-hGem); tetO-YAP S127A (kindly provided by Jonas Larsson^98^), which enables YAP1 activation in a doxycycline (DOX)-inducible manner; and Rosa26-rtTAM2 (stock #006965, Jackson Laboratory), with constitutive expression of a reverse tetracycline-controlled transactivator. Animals cohorts were used at exact post-natal days indicated, the adult cohort constitutes mice between 10-14 weeks of age, unless otherwise stated.

All strains were maintained in a C57BL/6 background. All experiments comprised a mixture of male and female mice, with no gender specific differences observed. For single cell RNA sequencing (scRNAseq) experiments exclusively, only male mice were used in order to avoid confounding effects due to estrous cycle.

### Esophageal tissue dimensions

Esophageal dimensions were measured from postnatal day (P) 2 up to the adult state (P70-onwards), when homeostasis is fully established^10^. C57/Bl6 wild type (wt) mice were culled and the esophagus collected at time points indicated. Tissues were excised, taking as reference two anatomical landmarks, from the most proximal part of the trachea (cricoid cartilage) to the most distal part of the esophagus (opening of the stomach fundus). Esophageal tubes were then cut open longitudinally and flattened in order to measure their length and width accurately under a dissecting microscope. Tissue area was then calculated using the obtained measurements. Tissue area averaged from n = 3-22 per time point.

### Histology

Hematoxylin and Eosin (H&E) staining was performed by the Histology Core Service at Cambridge Stem Cell Institute and imaged using Evos XL Core Cell Imaging System.

### Immunofluorescence

For wholemounting, esophagi were excised, cut open longitudinally, the muscle layer removed, and the tissue flattened under a dissecting microscope using fine forceps. The resulting epithelial-stromal composite was then fixed in 4% paraformaldehyde (Alfa Aesar; 043368) in PBS for 30 minutes. Epithelial-stromal composites were stained using an extended protocol due to their increased thickness. Samples were incubated for 30 minutes in permeabilization buffer (PB; 0.5% Bovine serum albumin (VWR International; 126575-10), 0.25% Fish skin gelatin (Sigma; G7765), 1% Triton X-100 (Fisher Scientific Ltd; 10102913) in PBS), and then blocked for 2 hours in PB containing 10% Donkey Serum (PBDS; Scientific Laboratory Supplies; D9663). Primary antibodies were incubated in PBDS for 3 days at 4°C followed by 4 washes over the next 24 hours with 0.2% Tween-20 (Promega UK Ltd; H5151) in PBS. Secondary antibodies were incubated overnight at 4°C in PBDS and were subsequently washed 4 times over 4 hours with 0.2% Tween-20 in PBS. Antibody details provided in **Supplementary Table 1**. To stain cell nuclei, a final incubation with 1 μg/ml DAPI (Simga; D9542) in PBS at 4°C was used before mounting the tissue. Wholemounts were cleared in 1.52 Rapiclear mounting media (SUNJin Lab; RC152001) for imaging. Organ culture samples were stained following the same procedure.

Wholemounts of the esophageal epithelium (EE) were used for staining with specific antibodies as indicated in figures. EE wholemounts were prepared by incubating esophagi in 5 mM EDTA (Life Technologies; 15575020) in PBS for 2.5 hours at 37°C following the removal of the muscle layer. Following incubation, the epithelium was gently peeled from the stroma using fine forceps. These layers were individually flattened and fixed as above. Staining was achieved by permeabilizing in low Triton PB (pB; 0.5% Bovine serum albumin, 0.25% Fish skin gelatine, 0.5% Triton X-100 in PBS) for 10 minutes, followed by blocking for 1 hour in pB containing 10% Donkey Serum (pBDS). Primary antibodies were incubated in pBDS overnight at 4°C and washed 4 times over 2 hours with 0.2% Tween-20 in PBS. Secondary antibodies were incubated at room temperature for 3 hours in pBDS before being washed again as with primary. Staining of cell nuclei was carried out as above with 1 μg/ml DAPI in PBS before mounting.

For staining of 10 μm tissue cryosections, fixed esophagi were embedded in optimal cutting temperature compound (OCT; Fisher Scientific Ltd; 12678646) for cryosectioning onto glass slides. Sections were permeabilized in pB for 10 minutes and then blocked in pBDS for a further 30 minutes prior to overnight incubation with primary antibodies in pB at 4°C. This was followed by a 1 hour incubation with secondary antibodies in pB at room temperature. Sections were washed 4 times for 5 minutes in 0.2% Tween-20 in PBS between incubations.

Where indicated, EdU incorporation was detected using Click-iT imaging kits according to the manufacturer’s instructions (Invitrogen). Figures show representative images of a minimum of 3 animals.

### EdU tracing

For *in vivo* tracing, EdU (Life Technologies; A10044) was administered 10ul/g body-weight at 0.1mg/ml via subcutaneous (s/c) injection at time points indicated, throughout postnatal development. Tissues were collected 2 and 24 hours after injection. For EdU tracing in organ cultures, EdU was added to the culture media at a 10 μM final concentration, and incubated for 2 hours at 37°C and 5% CO_2_. EdU positive basal and suprabasal cells were quantified in esophageal wholemounts from a minimum of 3 z-stack images per animal, from 3 animals, 11 basal cells analyzed. Quantifications were normalized by the number of basal DAPI+ cells per field where indicated.

To enable the progeny of single EdU+ cells to be tracked in adult mice, EdU was administered at 18:00 (s/c), when the proportion of S phase cells is low in the circadian cycle, making clonal density labelling feasible^31, 99^. 12 hours after the injection, individual post-mitotic EdU+ pairs were visualized by confocal microscopy in esophageal wholemounts. KLF4 expression profile was determined in 225 EdU pairs from 3 different animals.

### Basal cell density

Basal cell density was calculated at different postnatal time points by imaging the basal layer of esophageal wholemounts, and quantifying the number of DAPI+ basal cells per field in 3-6 random images per animal, from a minimum of 3 different mice.

### Total cell production and imbalance

Total cell production rate was calculated by first estimating the total number of basal cells in the whole esophagus at time points throughout PND. This was achieved by multiplying the basal cell density (defined as the number of DAPI+ basal cells per area) by the total tissue area at each time point. The total basal cell production rate, *Ḃ*, is then estimated as the average change in basal cell number, *B*, divided by the time step.

The degree of fate imbalance, i.e. the degree to which basal cell fate is biased towards duplication over differentiation and cell loss, is estimated as (1/*λ*)*Ḃ/B*, where *λ* is the cell division rate. The division rate is determined from the results of short-term (2h) EdU incorporation assay and normalized by the reported division rate of basal cells in the homeostatic adult tissue^10^.

### Tissue thickness

Epithelial tissue thickness, including terminally differentiated and enucleated cells, was measured using side views of rendered confocal wholemount images at a range of time points throughout postnatal development. 18 individual measurements were made for each of 3 animals.

### Imaging, analysis and segmentation

Confocal images were acquired using either inverted Leica SP5 with standard laser configuration, SP8-X with white light laser or upright SP8-X with standard laser configuration. Typical confocal settings used included: 40X objective, optimal pinhole, scan speed 400 Hz, line average 3, optimal step size, and resolutions of 512×512 or 1024×1024 pixels. For quantification, images where acquired with a digital zoom of 2X or 3X.

Images were reconstructed from optical sections using Volocity 5.3.3 software (PerkinElmer) and ImageJ 1.51W. KLF4, EdU, Fucci2a (mCherry and mVenus), as well as YAP intensity measurements were performed on Volocity 5.3.3 using semi-automated image segmentation. Intensity analysis were set to include cells above the 80^th^ percentile of KLF4 intensity (**Fig. 2f,g and Extended Data Fig. 2d,e**), which was visually confirmed as an optimal threshold for KLF4 bright cells (KLF4+) by experimenter. EdU was set as >0.2 as detected in representative confocal images.

Collagen fibers were imaged using second harmonic generation (SHG), without the need for immunostaining, alongside NucRed^647^ (nuclear staining; Life Technologies; R37106) in wholemount samples at key time points throughout postnatal development. Imaging was performed using an inverted Leica TCS SP5 acousto-optic beam splitter (AOBS) multiphoton laser scanning microscope equipped with a Modelocked Ti:Sapphire Laser (Chameleon Ultra II, Coherent Inc., 680-1080nm tuning range, 80-MHz repetition rate, 140-fs pulse width) that allows for SHG imaging. For image acquisition the excitation laser (Ti:Sapphire) was tuned to 900 nm, subsequently producing emission of SHG signal at 450 nm. Typical settings included: 40X objective, fully-open pinhole (600 μm), scan speed 400 Hz, line average 2, 0.15 μm step size, and resolution 1024 x 1024 pixels. A minimum of 3 images per animal (n=3) were captured, and the collagen fiber orientation was assessed using method outlined in **Supplemental Information1:** Quantitative Image Analysis.

For single cell image segmentation, single 2 μm z slices of the basal layer were used. Images were acquired using a Leica SP5 confocal microscope system. Confocal settings included: 40x objective; 3x digital zoom; line average 3; scan speed 400 Hz; resolution of 512×512 pixels. A minimum of 3 images were captured for each of 3 animals at indicated time points. Analysis utilized a deep-learning based image segmentation pipeline described in **Supplemental Information 1:** Quantitative Image Analysis.

### RNA sequencing

#### Single cell and RNA isolation

Epithelial-stromal composites from P7, P28 and Adult mice were obtained as indicated in “Immunofluorescence” section. For each sample, esophagi were pooled from different animals to enable the capture of sufficient cell numbers for sorting (5 mice for P7, 4 mice for P28 and 3 mice for Adult). Each time point included 3-6 independent samples.

Following initial preparation, esophageal whomeounts were cut into 3 mm^2^ pieces, and incubated in 0.5mg/ml Dispase (Sigma; D4818) in PBS containing 1U/μl RNAse Inhibitor (Life Technologies; AM2696) at 37°C for 10 minutes. Following incubation, the epithelium was carefully peeled away from the stroma using fine forceps and thoroughly minced before placing back into Dispase for a further 5 minute incubation at 37°C to create an epithelial cell suspension. EDTA was then added to the samples at a final concentration of 5 mM, and suspension diluted 1/5 by adding FACS staining buffer (SB; 2% heat-inactivated Fetal bovine serum (Life Technologies; 26140079), 25 mM HEPES (Life Technologies; 15630056), 1 mM EDTA in PBS) in order reduce Dispase activity. Single-cell suspension was obtained by filtering through a 30 μm cell strainer. Cells were then centrifuged at 300xg for 10 minutes at 4°C, and resuspended in SB. Staining with primary antibodies or isotype controls was performed for 15 minutes at 4°C, followed by a final wash in FACS buffer (2% heat-inactivated Fetal bovine serum and 25 mM HEPES in PBS) at 300 g for 5 minutes at 4°C. The final cell suspension was placed in FACS buffer containing 1U/μl RNAse Inhibitor.

The following immunoglobulins were used to enrich for the epithelial cell population in the esophagus: Ep-Cam-AF647 at a concentration of 20μg/ml to stain epithelial cells, and CD45-APC-Cy7 at a concentration of 8μg/ml to exclude immune contamination. Isotype controls were used at same concentration as specific antibodies: AF647 Rat IgG2a and APC-Cy7 Mouse IgG2a. Antibody details in **Supplementary Table 1**.

Please note, work was carried out using RNAse free and/or PCR grade sterile reagents and plastic ware wherever possible.

Stained samples were sorted on a BD FACSAria™ III cell sorter utilizing DAPI incorporation (4μg/ml) to exclude dead cells from sorted suspension. Viable epithelial (Ep-Cam^+^/CD45^-^/DAPI^-^) cells were sorted to produce samples of approximately 5,000 cells for downstream scRNAseq.

#### Library preparation and sequencing of RNA from single cells

scRNAseq libraries were generated using 10X Genomics kits (Single Cell 3’ v3) at the CRUK-CI Genomics Core Facility of the CRUK Cambridge Institute.. Libraries were generated in two different batches/dates. In one batch, 3 biological replicates for P7 and 3 for Adult were used for 6 libraries. In the second batch, 3 biological replicates for P28 and 3 for Adult were used for 6 libraries. Each biological replicate consisted of pooled cells from 5 mice at P7, 4 mice at P28 and 3 mice in Adult. The use of Adult samples in both batches provided common samples to control batch effect between the two. The cells for each biological replicate were loaded into one channel of a 10X Chromium microfluidics chip to package them into one library. The 12 libraries in total were sequenced on Illumina NovaSeq 2 SP and 2 S2 flow cells in the core facility above.

#### Single-cell RNA-seq analysis

##### Data processing for scRNAseq

The raw sequencing data from the 10X Genomics platform were processed using CellRanger (v3.0.2). CellRanger aligned reads, filtered empty dropouts and counted unique molecular identifiers (UMIs) to generate a count matrix. We used Ensembl GRCm38/mm10 (release 92) as the reference genome for the read alignment. To filter out low quality cells, cells, basic QC metrics were calculated using R package scater (1.12.2)^100^ and cells with less than 1251 genes were removed. In addition, cells with mitochondrial proportions above 15% were also discarded from further analysis. Genes expressed in less than 3 cells were removed. Read counts were normalized by a deconvolution method using the R package scran (v1.12.1)^101^. Cells expressing fibroblast marker genes including Col1a2, Col1a1, Fn1, and Pdgfra were considered as non-epithelial contaminants. To filter out these non-epithelial cells, the normalized read count values for Col1a2, Col1a1, Fn1, and Pdgfra were averaged and the cells with the averaged normalized values for the four genes more than 99 percentile (=0.4301828) of the distribution was filtered out as the contaminants.

##### Dimension reduction and data visualization

PCA combined with technical noise modeling was applied to the normalized data for dimension reduction, which was implemented by the denoise_PCA function in the R package scran. This denoise PCA does not strictly require explicit feature selection, such as highly variable genes. The data were then projected using two-dimensional Uniform Manifold Approximation and Projection (UMAP) or t-Distributed Stochastic Neighbor Embedding (t-SNE) with default parameter setting using the R package scater. The biological replicates for each of the three time points overlapped well with each other, confirming batch effects between different samples and conditions were negligible and batch effect correction was not necessary for further analysis. R package Seurat (v3.0.2)^102^ was used to visualize cells in the dimension reduction space.

##### Data clustering and cluster annotation

To perform clustering, we used a louvain community detection method^103^. First, a shared nearest-neighbor graph was constructed using k=20 nearest-neighbors of each cell (buildSNNGraph function in the R package scran). In this graph, two cells were connected by an edge if they shared nearest-neighbors, with the edge weight determined by the highest average rank of the shared neighbors. Then the Walktrap method from the R package igraph (v1.2.4.1) (with steps = 4 as the default option) was used to identify densely connected communities that were considered to be cell clusters.

Cell clusters were annotated based on differentially expressed genes and known marker genes for cell types. If a few neighboring clusters in the dimension reduction spaces shared key expression patterns, they were merged into one cell type manually. First, based on basal cell markers such as Krt14, Itgb1, and Itga6, and differentiation markers such as Tgm3, Krt13, and Grhl3, all clusters were classified into basal cells (clusters 1, 2, 3, 5, 6, 7, 10, 11, 12, 13, 15, 16, 17) and differentiated cells (clusters 4, 8, 9, 14) (**Extended Data Fig. 3b**).

Then we classified all cells into different cell cycle phases based on cell cycle genes: clusters 1, 2, 4, 6, 7, 8, 9, 12, and 14 for G0/G1; clusters 11 and 13 for G1/S; clusters 3, 5, 10, 15, 16, and 17 for S/G2/M (**Extended Data Fig. 3e,f**). As a result, all cells were annotated as one of three cell types as follows: clusters 3, 5, 10, 11, 13, 15, 16, and 17 for basal cycling cells; clusters 1, 2, 6, 7 and 12 for basal resting cells; clusters 4, 8, 9, and 14 for differentiated cells (**Extended Data Fig. 3c,d**). For cluster annotation, we visualized gene expression in the dimension reduction and in violin plots using built-in functions of Seurat (v3.0.2).

##### Classifying gene expression for basal cells into distinct dynamic patterns along the time course and Gene Ontology analysis of the different expression patterns

After annotating cell types, we then focused on molecular change for basal cells over time. To this end, we first identified differentially expressed (DE) genes for each time point using the function ‘findMarkers’ of scran (v.1.12.1) by defining them as the genes having FDR < 0.05 and absolute value of log2(fold-change) > (95 percentile of all log2(fold-change)) in the three comparisons of P7 vs. P28, P28 vs. Adult, P7 vs. Adult. The total number of DE genes was 1738. We then scaled the normalized expression value of DE genes for all cells and calculated the average of the scaled expression value across the cells for each time point. Then scaled log2(fold-change) for each time was calculated by subtracting the averaged scaled value at P7 from those at all time points (thus, scaled log2(fold-change) at P7 = 0).

The scaled log2(fold-change) profiles for DE genes (i.e. 1738 genes) were clustered based k-means clustering (k=10) using pheatmap (1.0.12) and the resulting 10 clusters were grouped into 4 major patterns (**Extended Data Fig. 3e,f**): 148 genes for Pattern 1; 545 genes for Pattern 2; 153 genes for Pattern 3; 892 genes for Pattern 4. The genes in each major pattern was used for Gene Ontology analysis by DAVID (https://david.ncifcrf.gov/)^104^. The resulting GO terms were manually curated and selected based on our observation in experiments and known biological knowledge (**Fig. 3f**).

### Blebbistatin treatment of adult esophageal organ culture

After opening the esophageal tube longitudinally, the muscle layer was removed with fine forceps, and cut using a scalpel into approximately 3×4 mm pieces. Tissue pieces were placed onto transparent ThinCert™ inserts (Greiner Bio-One Ltd; Cat# 657641), stroma side down to the insert membrane, leaving the epithelium facing upward. Tissues were allowed to dry for 5 minutes at 37°C to ensure attachment to the membrane, and cultured in minimal medium containing 1 part of DMEM (4.5 g/L D-Glucose, Pyruvate, L-Glutamine; Life Technologies; 41966029): 1 part DMEM/F12 (Life Technologies; 11320033), supplemented with 5 μg/ml insulin (Sigma; I5500), 1.8×10-4 M adenine (Sigma; A3159), 5% fetal calf serum (PAA Laboratories; A15-041), 5% Penicillin-Streptomycin (Sigma; P0781) and 5 μg/ml Apo-Transferrine (Sigma; T2036). After culture at 37°C, 5% CO_2_ for 3 hours to ensure adequate tissue adhesion to the insert membrane, 25 μM Blebbistatin (Cayman Chemicals; 13165) was added to the media of experimental explants. Control explants were treated with DMSO vehicle alone. Following a further 45 hours, explants were removed and fixed using 4% paraformaldehyde.

### Whole esophageal organ culture and tissue stretching

Design of the 3D printed stretcher was developed from prototype in^78^ using Fusion360 Autodesk and Ultimaker Cura software and printed using Ultimaker^3^ 3D printer in ABS (biocompatible plastic). Once printed and components sterilized, a 10 cm screw (adjustable screw) was inserted into the lower clamp to allow for fixing its placement and control tissue stretching (See **Fig. 7b**). The device had a millimetric scale printed to its left to guide the percentage of stretch applied to the tissue.

Once the sterile device was assembled, and tissue clamped to it at both ends, the system was placed in a sterile plastic container with lid. The set up allows for the adjustable screw to be accessible from the outside of the container in order to set up the tissue stretching conditions.

For *in vitro* stretching, EE samples were collected and the muscle layer removed by careful pulling with fine forceps whilst retaining the tubular structure. Samples were then placed into the upper and lower clamps under no tension. Clamps were tightened using M3 nuts and screws until samples were held in place at each end. The lower clamp could then be pulled back using the adjustable screw on the exterior of the device. The millimetric scale was used to apply a 40% stretch in tissue length. Control samples were only clamped at one end (top clamp, see **Fig. 7b;** inset) and cultured at their original length exposed to no longitudinal tension. Whole-organ cultures were kept for 48 hours at 37°C, 5% CO_2_ in minimal medium containing 1 part DMEM (4.5 g/L D-Glucose, Pyruvate, L-Glutamine; Invitrogen 11971-025): 1 part DMEM/F12, supplemented with 5 μg/ml insulin, 1.8supplemented with 5 10-4 M adenine, 5% fetal calf serum, 5% Penicillin-Streptomycin and 5 μg/ml Apo-Transferrine. For EdU incorporation assays, EdU was added to media at a final concentration of 10 μM, 2 hours prior to sample collection.

### Statistics and Reproducibility

Number of biological replicates and number of animals are indicated in figure legends. Data are expressed as median values ± SEM unless otherwise indicated.

Differences between groups were assessed by using two-tailed unpaired t-test, one-way or two-way analysis of variance (ANOVA) as indicated in figure legends. ANOVA based analysis was followed by Tukey’s test for multiple comparisons. Statistical differences between groups were assessed utilizing GraphPad Prism software. No statistical method was used to predetermine sample size. Experiments were performed without methods of randomization or blinding.

The methods used for deep learning based segmentation to study tissue spatial reorganization throughout postnatal development are set out in **Supplemental Information 1:** Quantitative Image Analysis.

## Data and Code Availability

All data and codes used and/or generated in this study are available upon request to the authors. Image segmentation and computational image analysis was performed using Python codes developed and/or adapted for this study. The statistical analysis of the single-cell RNA-seq data was performed using R codes developed for this study. The software will be made available upon request.

## Acknowledgments

We thank members of the Alcolea lab, Simons lab and Bartomeu Colom (Sanger Institute) for comments and suggestions; the staff of the University Biomedical Services, Gurdon Institute; Peter Humphreys in imaging core facilities at JCBC; Alex Sossick, Richard Butler and Nicola Lawrence at the Gurdon Institute Imaging Facility; Jonas Larsson (tetO-YAP S127A); Ian J. Jackson (Fucci2a); NIHR Cambridge BRC Cell Phenotyping Hub for FACS support; the Jeffrey Cheah Biomedical Centre (JCBC) core facilities. We thank T. Savin, E. Hannezo, B. Ladoux, G. Charras, S. Hénon, N. Harmand, G. Duclos and J-B. Lugagne for their insight on deep learning based image analysis and/or tissue mechanics. This work was mainly supported by funding from the Wellcome Trust and The Royal Society (105942/Z/14/Z to M.P.A), and a core support grant from the Wellcome and MRC to the Wellcome-Medical Research Council Cambridge Stem Cell Institute. J.M was supported by a CRUK Cambridge Cancer Centre PhD fellowship. The work also received support from the Novo Nordisk Foundation (NNF18CC0033666 to K.K; NNF17OC0028730 to K.B.J and NNF17CC0027852 to K.B.J); the Wellcome Trust (098357/Z/12/Z Junior Interdisciplinary Research Fellowship to A.H); Human Frontier Science Program (LT000092/2016-L to S.H); National Institutes of Health (ZIA BC 011763 to R.I.B); Wellcome PhD stutentship (to F.J.E); The Royal Society and European Research Council (‘CellFateTech’, 772798 to K.J.C). B.D.S. acknowledges funding from the Wellcome Trust (098357/Z/12/Z and 219478/Z/19/Z) and the Royal Society in the form of an E.P. Abraham Research Professorship (RP\R1\180165).

## Author Contributions

J.M, F.J.E, R.I.B designed, validated and conducted experiments with the support from M.P.A; K.K and S.U carried out experiments under K.B.J supervision; A.H developed deep learning segmentation imaging pipeline, and performed associated tissue spatial reorganization and SHG analysis; S.H guided the experimental design for single-cell RNA sequencing and performed analysis of the data. A.H and S.H were both under B.D.S supervision; C.V designed and provided expertise on the 3D printed stretcher device with K.J.C input; K.J.C, R.I.B and A.H offered advice and technical expertise on tissue mechanic aspects; K.B.J and B.D.S supervised parts of the study and provided expertise in the epithelial stem cell field; J.M and M.P.A conceived the project, supervised experiments, and wrote the original manuscript draft with input from B.D.S. Review & Editing of final manuscript by all authors. Funding acquisition by A.H, R.I.B, K.B.J, D.B.S, and M.P.A.

