## Supplementary file for "A biomechanical switch regulates the transition towards homeostasis in esophageal epithelium"

**Extended Data Figures and Legends.**

**Extended Data Fig. 1:** Related to Fig. 1

**Extended Data Fig. 2:** Related to Fig. 2

**Extended Data Fig. 3:** Related to Fig. 3

**Extended Data Fig. 4:** Related to Fig. 3

**Extended Data Fig. 5:** Related to Fig. 4

**Extended Data Fig. 6:** Related to Fig. 5

**Extended Data Fig. 7:** Related to Fig. 7

**Supplementary Information**

**Supplemental Information:** Quantitative Image Analysis

**Supplementary Table 1.** Antibodies list

### Extended Data Fig. 1: Postnatal characterization

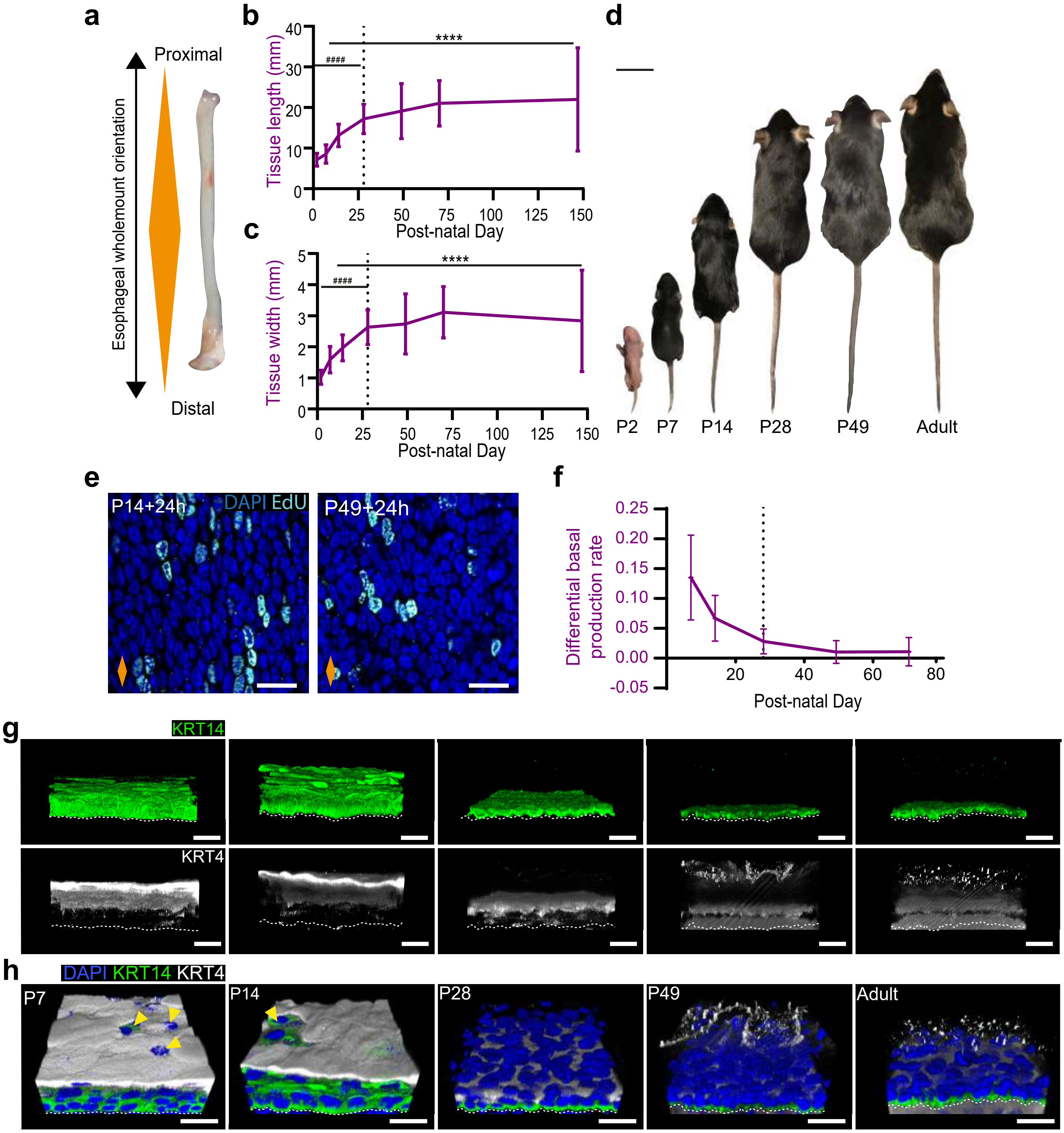

**Extended Data Fig. 1: Postnatal characterization. Related to Fig. 1.**

**a,** Diagram illustrating longitudinal esophageal orientation from proximal to distal as marked by orange diamond. **b,** and **c,** Esophageal tissue growth in length and width over time, respectively. **d,** Images showing animal body growth throughout postnatal development. **e,** Representative images showing EdU+ basal cells 24 hours post-labelling in P14 and P49 from **Fig. 1e-g**. **f,** Graphical representation of differential basal production rate throughout postnatal development. See Methods. **g,** 3D rendered z-stacks showing split confocal channels from **Fig. 1i**. **h,** Typical 3D rendered confocal z-stacks showing tilted side views from **Fig. 1i**. Yellow arrows indicate immature epithelial barrier. **Scale bars**. S1D(2 cm); S1E,G-H(20 µm). **Stainings**. Blue, DAPI; cyan, EdU; green, KRT14; greyscale, KRT4. Dashed white lines indicate basement membrane.

All data derived from wild-type *C57BL/6J* mice, expressed as mean ± SEM and analyzed using one-way ANOVA with Tukey's multiple comparisons test (n = 103; $^{\#\#\#\#}p < 0.0001$ relative to P70; $^{****}p < 0.0001$ relative to P7).

Dashed lines indicate P28. Orange diamonds depict longitudinal orientation of the esophagus where indicated.

### Extended Data Fig. 2: KLF4 basal cell prolife in FUCCI2a mice

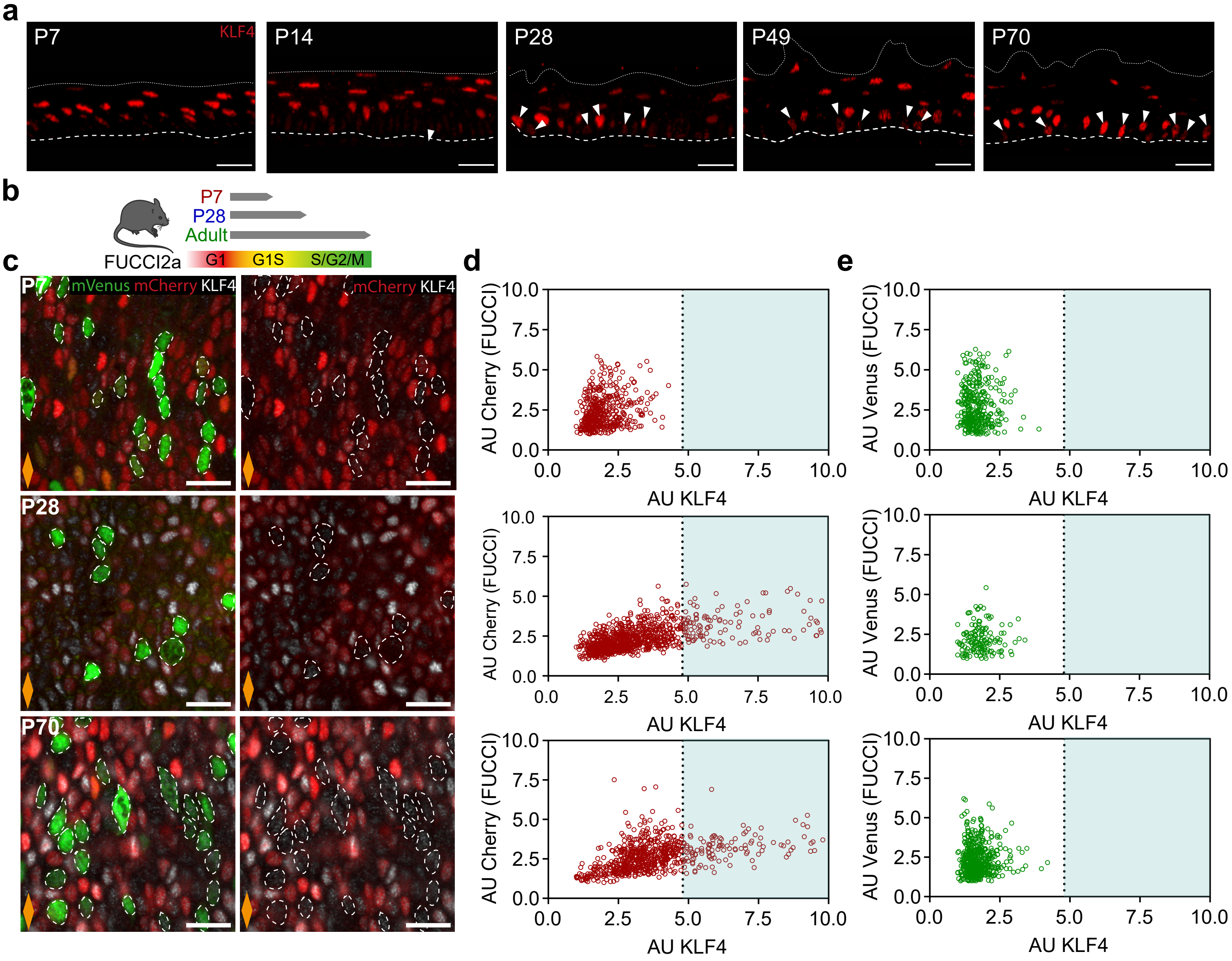

**Extended Data Fig. 2: KLF4 basal cell prolife in FUCCI2a mice. Related to Fig. 2.**

**a,** Representative confocal z-stacks showing side views of EE wholemounts from **Fig. 2a**. Dashed lines indicate basement membrane; dotted lines mark the upper limit of the EE. Red, KLF4. **b,** *In vivo* protocol. Esophagi from FUCCI2a mice were collected at time points indicated. Schematic indicating expression pattern of fluorescent proteins in FUCCI2a mouse model. **c,** Confocal images showing basal views of typical FUCCI2a EE wholemounts in **(b)**. Orange diamonds indicate longitudinal orientation of the esophagus. Green, mVenus; red, mCherry. **d** and **e,** Correlation between KLF4 protein expression and reporter fluorescent proteins mCherry **(d)**/mVenus **(e)** in the basal layer from **(b)** and **(c)**. **Scale bars** 20 µm.

### Extended Data Fig. 3: Single cell RNA sequencing annotation

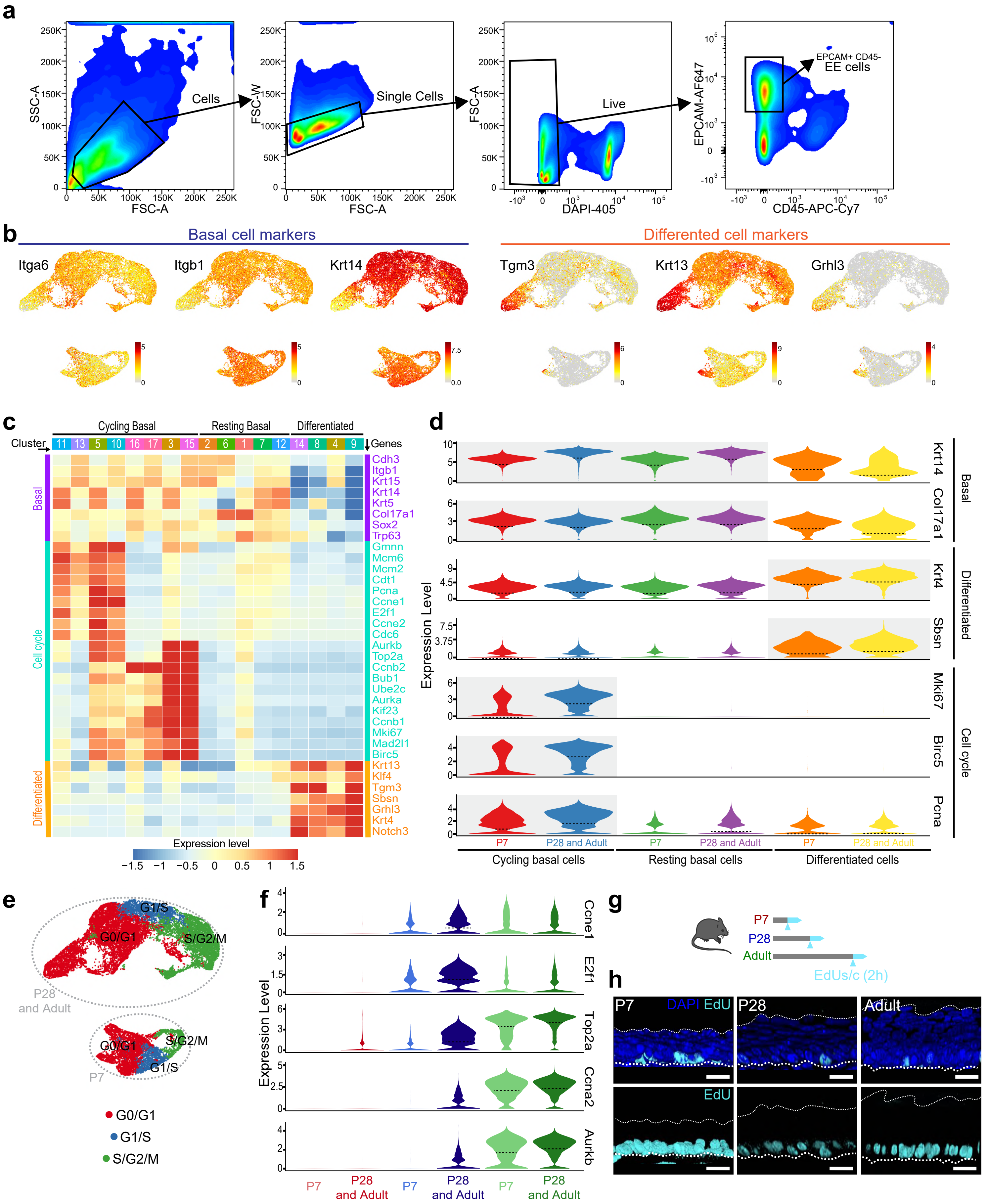

**Extended Data Fig. 3: Single cell RNA sequencing annotation. Related to Fig. 3.**

**a,** Flow cytometry gating strategy for isolation of EE cells (EpCam+/CD45-). Representative plots from adult sample. 3-6 samples per time point were analyzed. **b,** UMAPs showing expression of representative makers for basal (left panels) and differentiated cells (right panels) in EE. **c,** Heatmap showing expression of representative marker genes for basal cells, cell cycle, and differentiation for the 17 clusters shown in **Fig**. **3c** (cluster number in upper bar). The expression values were log2-transformed normalized UMIs followed by scaling and averaging across cells in the same clusters. **d,** Violin plots showing expression of representative epidermal (basal vs. differentiated) and cell cycle markers at different postnatal stages (P7 vs. P28 and Adult) split by annotated cell cohorts in **Fig. 3d**. **e,** UMAP showing spatial distribution of distinct cell cycle phases. The cell cycle phases were annotated using cell cycle analysis by R package scran (v 1.12.1) combined with manual curation based on expression of cell cycle genes in **(c)**. **f,** Violin plots showing expression of representative cell cycle genes at postnatal stages split by cell cycle cohorts identified in **(e).** The color scheme for cell cycle phases is the same as that in **(e)**. **g,** *In vivo* protocol. Mice were treated with a single EdU injection 2 hours prior culling at the time points indicated. **h,** Typical 3D rendered confocal z-stacks showing side views of EE wholemounts. Dashed white lines, basement membrane. Dotted white lines, upper EE limit. Blue, DAPI; cyan, EdU; scale bar 10 µm. For violin plots in **(d)** and **(f)** expression level means log2-transformed normalized UMIs and dotted lines indicate the median of the distribution. Color bars of UMAPs in **(b)** indicate log2-transformed normalized UMIs.

### Extended Data Fig. 4: Single cell RNA sequencing expression profile

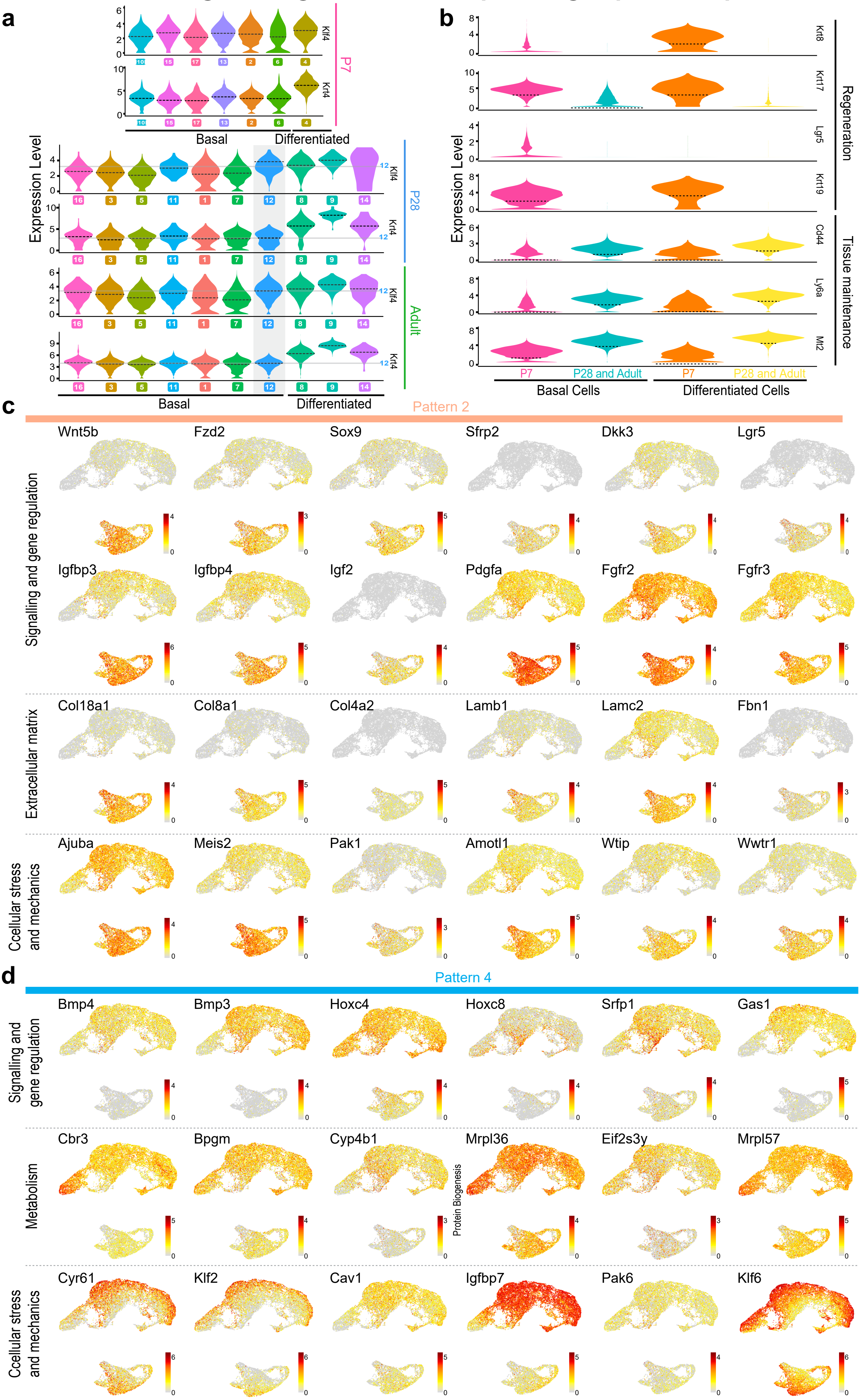

**Extended Data Fig. 4: Single cell RNA sequencing expression profile. Related to Fig. 3.**

**a,** Violin plots showing expression of Klf4 and Krt4 for cells in individual clusters at P7 (upper), P28 (middle) and Adult (lower). **b,** Violin plots showing expression of genes associated with regeneration vs. homeostasis for different epithelial cell types (Basal vs. Differentiated) at distinct postnatal stages (P7 vs. P28+Adult). Basal cells include both cycling and resting cells from **Fig. 3d**. **c** and **d,** UMAP showing expression of genes related to key biological processes from Gene Ontology analysis for Patterns 2 **(c)** and 4 **(d)** in **Fig. 3e,f**. For violin plots in **(a)**, and **(b)**, expression level means log2-transformed normalized UMIs and dotted lines indicate the median of the distribution. Color bars of UMAPs in **(c)** and **(d)** indicate log2-transformed normalized UMIs.

#### Extended Data Fig. 5: Deep Learning based segmentation

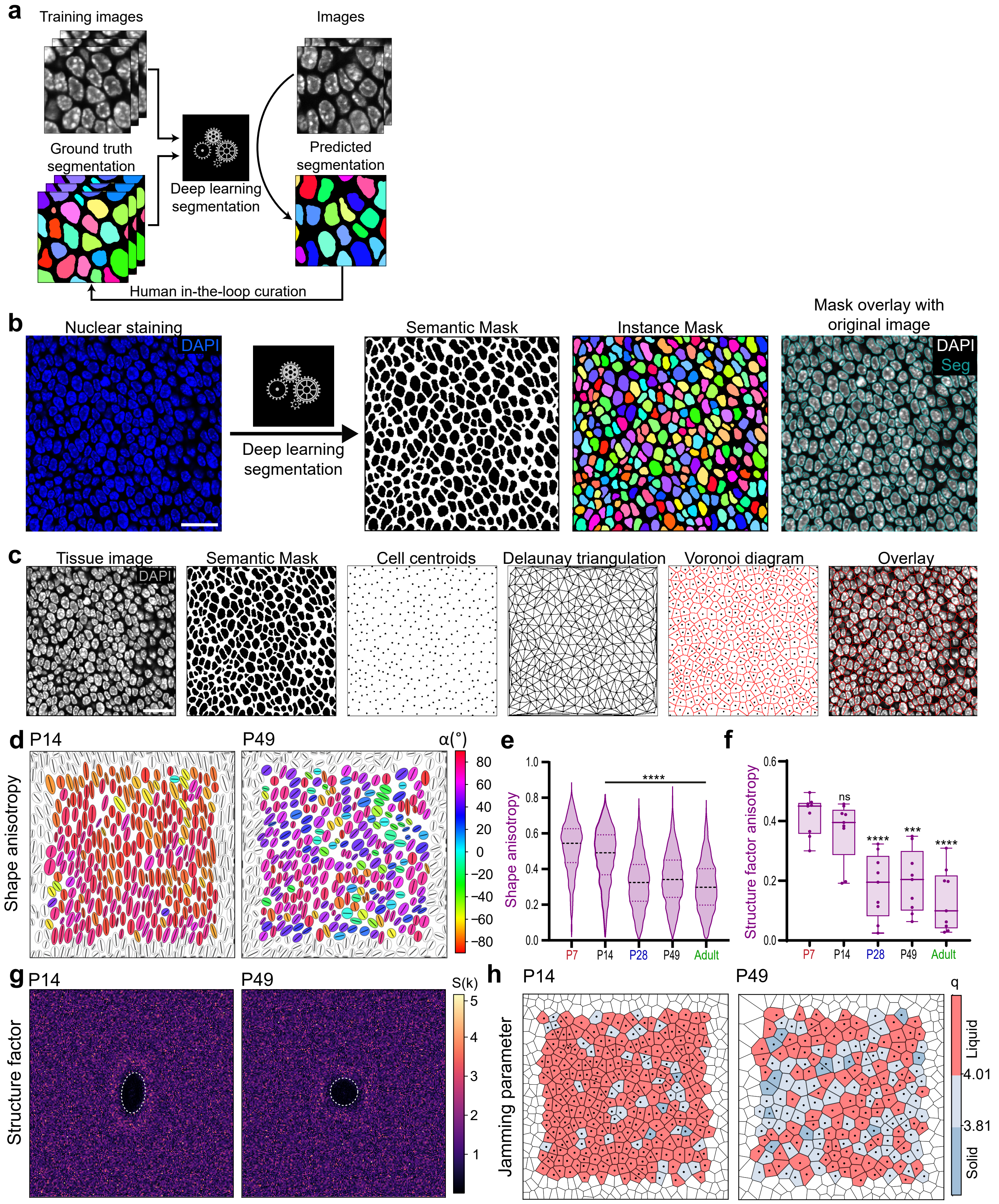

**Extended Data Fig. 5: Deep Learning based segmentation. Related to Fig. 4.**

**a,** Schematic depicting the principle of deep learning based segmentation. Manually or semi-automatically annotated “ground truth” images of the tissue are used to train a U-Net convolutional neural network to produce an automated segmentation mask of cellular features, such as the cell nuclei. Training of the network is assessed on a set of “validation” images and iteratively optimized until satisfactory segmentation performance is achieved. **b,** Schematic of pipeline utilized for the segmentation on single z-slice confocal images of the EE basal layer. Nuclear segmentation was based on DAPI staining (blue). Mask panels show the semantic (binary) and instance segmentation masks obtained as outputs of the pipeline. Mask overlay shows the match between the binary mask and the original fluorescence image. Scale bar, 20 µm. **c,** Schematic describing the computation of Voronoi diagrams of the tissue. Single z-slice confocal images of the EE basal layer are segmented using the pipeline described in **(b).** Cell centroids are computed using the obtained binary mask. Delaunay triangulation of cells in the images is performed using the coordinates of cell centroids. Voronoi diagrams are calculated as the dual of Delaunay triangulation of cells in the tissue and overlayed onto the original fluorecence image. Scale bar 20 µm. **d,** Cell shape anisotropy tensor represented as an ellipse calculated from the nuclear centroid position of each basal cell at P14 and P49 (supplementary to **Fig. 4d**). Long axis of each ellipse is proportional to the dominant eigen value of the tensor. Orientation is color-coded. Results from a representative experiment are shown; n=3. **e,** Violin plots showing the distribution in cell shape anisotropy throughout postnatal development. n=2052-2594 number of segmented cells from 3 animals per time point. Black dashed line, median. One-way ANOVA with Tukey’s multiple comparisons test (****$p < 0.0001$ relative to P7). **f,** Structure factor shape anisotropy distribution as shown in **(g)** and **Fig. 4e**. Data represented as box plot; individual measurements from n=3. **g,** Bidimensional structure factor quantifying basal cell spatial organization at P14 and P49 (supplementary to **Fig. 4e**). Changes in the dashed white outline (from ellipse to circle) depict a transition from anisotropic to isotropic cell distribution over time; n=3. **h,** Jamming parameter ($q=P/\sqrt{A}$) represented as a voronoi diagram calculated from the centroid of each cell at P14 and P49 (supplementary to **Fig. 4g**). Blue, solid-like “jammed”state; red, liquid-like state. Results from a representative experiment are shown; n=3.

All data derived from wild-type *C57BL/6J* mice.

### Extended Data Fig. 6: Esophageal tissue strain and Second harmonic generation

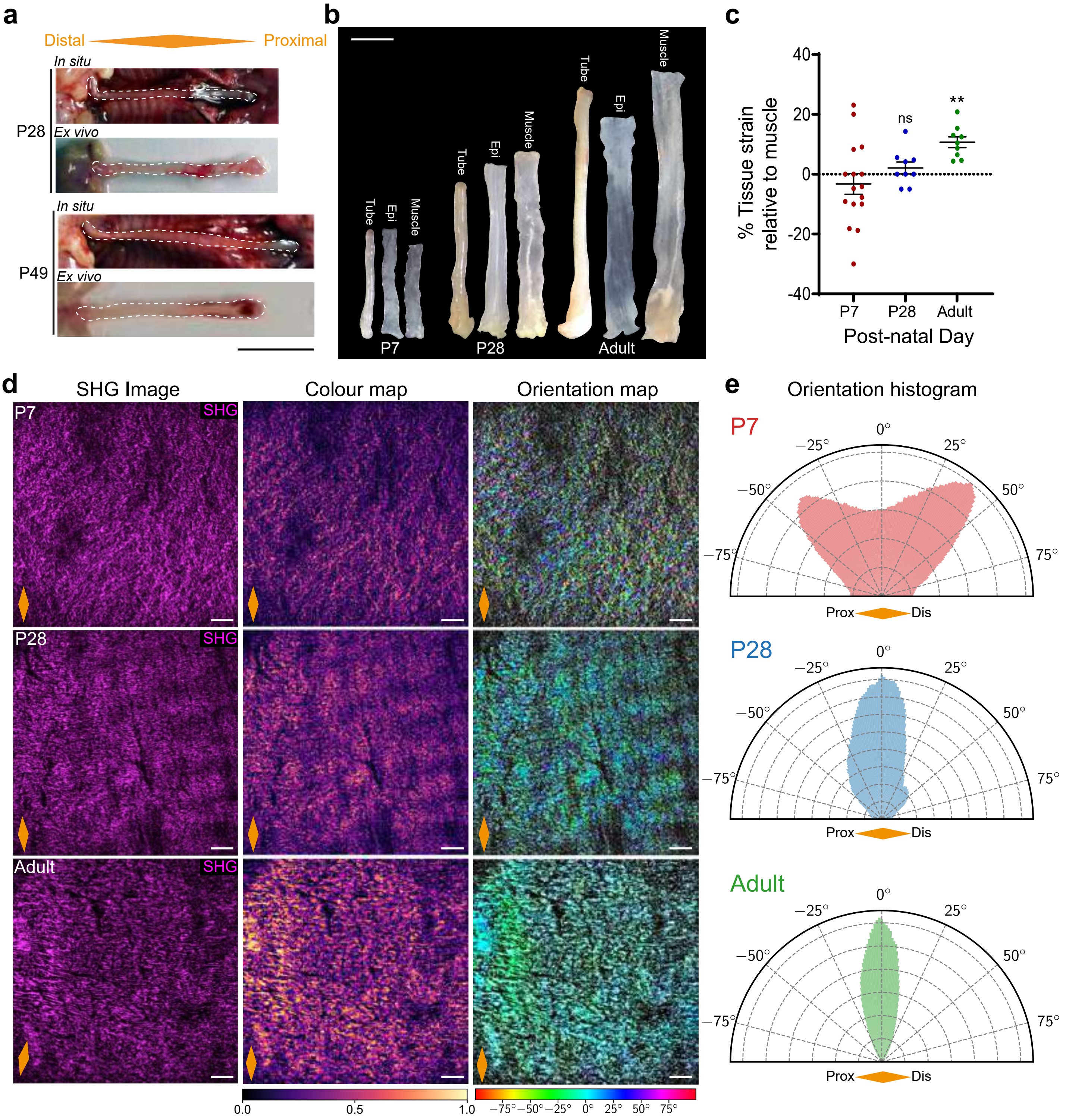

**Extended Data Fig. 6: Esophageal tissue strain and Second harmonic generation. Related to Fig. 5.**

**a,** *In situ* and immediate *ex vivo* images of esophageal tubes at P28 and P49 (supplementary to **Fig. 5b**). White dashed lines delineate esophageal tube; scale bar 1 cm. **b,** Representative images showing the size of combined and separate esophageal layers; Full esophageal tube (Tube), epithelial composite (Epi) and muscle layer (Muscle); scale bar 5 mm. **c,** Longitudinal tissue strain relative to muscle, represented as percentage. Data expressed as mean ± SEM. One-way ANOVA with Tukey's multiple comparisons test (n=6-9; $^{**}p < 0.01$; ns, not significant; relative to P7). **d,** Representative views of stroma underlying EE basement membrane using second harmonic generation (SHG). Left panels, collagen in magenta. Middle panels, color map of SHG signal intensity. Right panels, color-coded local orientation map of SHG signal. Scale bar 100 µm. **e,** Representative histograms depicting orientation distribution of collagen fibers in **(d)**. n=3.

All data derived from wild-type *C57BL/6J* mice. Orange diamonds depict longitudinal orientation of the esophagus where indicated.

### Extended Data Fig. 7: EdU incorporation assays in organ cultures

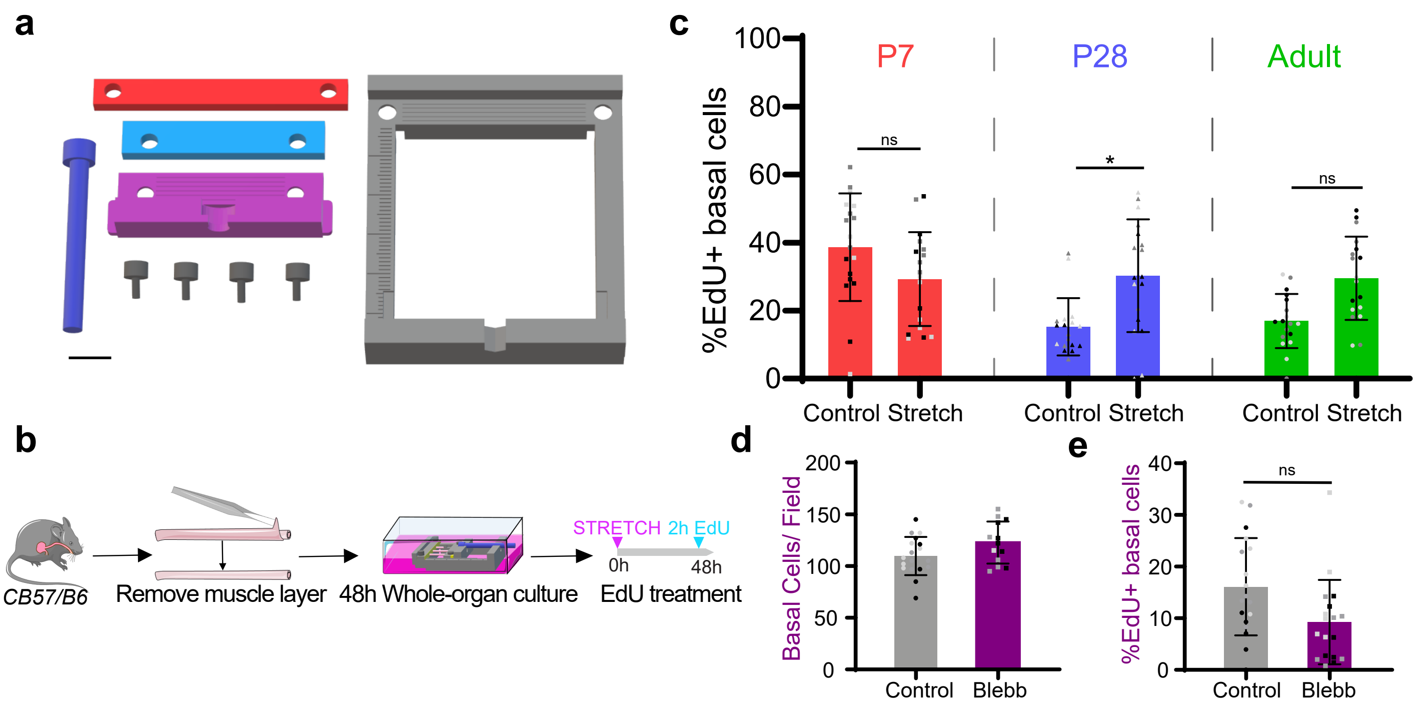

**Extended Data Fig. 7: EdU incorporation assays in organ cultures. Related to Fig. 7.**

**a,** 3D model of individual parts required for 3D printing of stretcher device. Scale bar 1 cm. **b,** *In vitro* protocol. Esophagi were collected and the muscle removed whilst maintaining tubular structure. Tissues were exposed to a 40% stretch using 3D printed stretcher and kept *in vitro* as whole-organ cultures for 48 hours. EdU was added to the media 2 hours prior collection. **c,** Basal quantification of EdU+ cells expressed as percentage of DAPI+ cells from **(b)**. n=3. **d,** Basal cell density, expressed in number of cells per field after BLEBB treatment in **Fig. 7e**. n=3. **e,** Basal quantification of EdU+ cells expressed as percentage of DAPI+ cells after BLEBB treatment in **Fig. 7e**. n=3.

All data are presented as mean ± SD and were analyzed using Unpaired t test (*p < 0.05; ns, not significant). Individual points show individual measurements, greyscale indicates values from each of 3 mice.

**Supplemental Information.**

**Quantitative Image Analysis**

**1. Tissue image segmentation**

Prior to analysis, all microscopy images were converted into 8 bit TIF files from their native proprietary file format. In addition to immunostainings of interest, DAPI was always used to label nuclei in wholemounts of the esophageal epithelium (EE). Nuclei were then segmented using an automated deep learning based image segmentation framework coded in Python (using Keras and TensorFlow libraries) and described in greater details elsewhere (A. Hallou, unpublished).

For each experiment, a "training" image set of six representative images was selected for each time point, with images taken from at least three biological replicates. A "ground truth" segmentation mask was obtained for each image of the training set using a semi-automated pipeline based on "ilastik", a random forest classifier based machine learning library for image segmentation [1]. For each condition, the random forest classifier was trained to distinguish between background and foreground pixels of interest producing a probability map of the image of interest, giving each pixel a particular probability to belong to a given class (background or foreground). A binary mask of the image was then obtained using a thresholding of the probability map, followed by post-processing steps composed of morphological opening/closing operations, size filtering and manual corrections.

To achieve full automation of image segmentation, we adopted a deep learning approach based on a U-Net network, a family of convolutional neural networks with an encoder/decoder architecture recently used for a variety of computer vision tasks, including biological and medical image analysis [2]. As shown in **Extended Data Fig. 4a**, we trained our U-Net network for each experiment using the training image set and associated ground truth binary masks prepared at the previous step. Training images were first subdivided into two groups, a first group containing two-thirds of the images was used to train the network while a second group, composed of the remaining images, was used for training validation.

Training was performed at original image resolution and image intensity was normalised between 0 and 1. Random rotation and elastic deformation were used as image augmentation techniques. The "loss function" used for training was pixel-weighted soft-max cross-entropy, with weighting used to enforce instance separation and class-balance (background vs. foreground objects). For initial training, the loss function was minimised on the training set using a stochastic gradient descent algorithm at a basal learning rate of $10^{-3}$ with a momentum 1 of 0.9 and momentum 2 of 0.999 for around 2.0-3.0 x $10^5$ time-steps up until reaching a learning rate decay of less than $10^{-4}$ for more than $10^4$ time-steps. Performance of the training was assessed on validation images using intersection over union (IoU) as a main metric for semantic segmentation accuracy, and was iteratively optimised until an IoU>93±3% was achieved for all training image sets.

Once the network was trained, other images of the same experiment were processed automatically to obtain binary nuclear masks as illustrated in **Extended Data Fig. 4b**. Instance objects, i.e. individual nuclei, were subsequently identified using a seeded watershed algorithm, using the ultimate points of binary mask as seeds [3].

#### 2. Spatial organisation and morphological properties of cell extraction and analysis

Using the instance mask of each image, 20 different individual spatial and morphological properties of each nucleus were systematically extracted: 1, surface area; 2, centroid x position; 3, centroid y position; 4, perimeter; 5, bounding box centre x position; 6, bounding box centre y position; 7, bounding box width; 8, bounding box height; 9, ellipse major axis; 10, ellipse minor axis; 11, ellipse major axis angle (with image x axis); 12, circularity; 13, Feret diameter; 14, Feret x start position; 15, Feret y start position; 16, Feret angle (with image x axis); 17, minimum Feret diameter; 18, aspect ratio; 19, Roundness and 20, Solidity. Generated data were stored in individual CSV files and subsequently analysed using custom Python scripts.

The point configuration made by the ensemble of nuclear centroids was used as the start point for further analysis of the tissue spatial organisation as shown in in **Extended Data Fig. 4c**.

The basal cell density, $\rho$, was evaluated as the average number of centroid points, $N$, per surface area, $L^2$, in 2D single-plane images of the EE basal layer:

$$\rho = \left\langle \frac{N}{L^2} \right\rangle$$

where $\langle .. \rangle$ denotes an average over all cells in the image. From this result, we obtained the characteristic neighbour distance between basal cells,

$$l_c = \frac{1}{\sqrt{\rho}}$$

To further quantify the spatial organisation of tissue, the bidimensional structure factor, $S(\boldsymbol{k})$ was derived, which characterises the long-range spatial organisation of the tissue:

$$S(\boldsymbol{k}) = \frac{1}{N} \left| \sum_{j=1}^{N} exp\left(i\boldsymbol{k} \cdot \boldsymbol{r}_j\right) \right|^2$$

Here $\boldsymbol{r}_j$ denote centroid position vectors of nuclei and $\boldsymbol{k} = \binom{k_x}{k_y}$ is the wave vector [4]. For practical purposes, the non-physical "forward scattering" contribution ($\boldsymbol{k} = 0$) was excluded, and $\boldsymbol{k}$ was computed using periodic boundary conditions, which is a reasonable assumption given that $L$ is much smaller than the overall tissue size:

$$k_x = k_y = \frac{2\pi n}{L} (n \in \mathbb{N}).$$

The values taken by the structure factor at small wave numbers reflect the degree to which there exists a large-scale spatial organisation of cells in the tissue and thus, if the tissue possesses positional order, $S(\boldsymbol{k})$ should significantly deviate from unity when **k** is small [5]. The structure factor anisotropy was then derived to quantify the extent to which the tissue spatial organisation varies along the different spatial coordinates [6]. To do so, the bidimensional structure factor was thresholded such that only values well below unity were kept:

$$\bar{S}(\boldsymbol{k}) = S(\boldsymbol{k}) \preccurlyeq 1.0.$$

The shape anisotropy tensor of the thresholded structure factor, $\bar{S}(\boldsymbol{k})$, could then be derived as:

$$\boldsymbol{I}_{\bar{S}} = \begin{pmatrix} \langle k_x k_x \rangle & \langle k_x k_y \rangle \\ \langle k_x k_y \rangle & \langle k_y k_y \rangle \end{pmatrix}$$

where $k_x$ and $k_y$ are now the coordinates of all the points of the thresholded structure factor with their origin taken at their barycentre $\langle k_x \rangle = \langle k_x \rangle = 0$, with brackets indicating an average over space. The four terms that appear in $\boldsymbol{I}_{\bar{S}}$ represent the coordinate covariances. From the eigenvalues of $\boldsymbol{I}_{\bar{S}}$, $\lambda_1$and $\lambda_2$, one can obtain the anisotropy of the structure factor as:

$$A_{\text{SF}} = \frac{\lambda_1 - \lambda_2}{\sqrt{\lambda_1^2 + \lambda_2^2}}.$$

From the coordinates of nuclei centroids, it is also possible to retrieve Voronoi diagrams of the epithelial tissue as the dual of their Delaunay triangulation [7]. We used the standard Delaunay triangulation algorithm provided by the SciPy library, but calculated modified Voronoi diagrams to handle infinite cells at the boundary of the image as shown in **Extended Data Fig. 4c**. Voronoi diagrams were used to quantify the local topology of the tissue and also as a convenient way to display various measured quantities that were colour-mapped onto them.

It is possible to carry out a similar analysis of the spatial organisation of the tissue at the individual cell level. Indeed, from the knowledge of the coordinates of every segmented cell outline, it was possible to derive the cellular shape anisotropy tensor:

$$\boldsymbol{I_c} = \begin{pmatrix} \langle xx \rangle & \langle xy \rangle \\ \langle xy \rangle & \langle yy \rangle \end{pmatrix}$$

where $x$ and $y$ are now the coordinates of all the points of the cell contour with their origin taken at their barycentre $\langle x \rangle = \langle y \rangle = 0$ (i.e. at the cell centroid), with brackets indicating an average over space. From the eigenvalues of ***I***, $\lambda_1$and $\lambda_2$, we can obtain the cell shape anisotropy $A_{\text{c}}$:

$$A_{\text{c}} = \frac{\lambda_1 - \lambda_2}{\sqrt{\lambda_1^2 + \lambda_2^2}}$$

The determination of the eigenvectors, allowed determination of the orientation of the long axis,$\theta$, and short axis, which is perpendicular to it. Also, knowing $\lambda_1$, $\lambda_2$ and $\theta$, one can represent the shape anisotropy tensor of each cell as an ellipse centred at its centroid and whose orientation is given by $\theta$, and long and short axis lengths by $\lambda_1$and $\lambda_2$ respectively.

Following the Landau-De Gennes theory of liquid crystals, the distribution of orientation of the long axis of each cell can also be used to quantify the orientational order of cells in the tissue, defining an orientation tensor [8]:

$$\boldsymbol{O} = \begin{pmatrix} \langle \cos\theta^{\,2} \rangle & \langle \cos\theta \sin\theta \rangle \\ \langle \cos\theta \sin\theta \rangle & \langle \sin\theta^{\,2} \rangle \end{pmatrix}$$

Basic linear algebra allows determination of the dominant eigenvalue of $\boldsymbol{O}$**,** which is called the nematic order parameter,

$$S = \sqrt{\langle \cos\theta^{\,2} \rangle + \langle \sin\theta^{\,2} \rangle}$$

This quantity is averaged over all segmented cells in an image and is thus a readout of the average orientational order in the tissue. The limiting case *S=0* would describe a completely disordered tissue, where cellular orientations are uncorrelated, while the limiting case *S=1*, would describe a perfect nematic order where all cells align with their neighbours [9].

The shape of each cell in the epithelium was further characterized by the "jamming parameter" or cell shape index, $q_j$, which is defined as the ratio of the cell perimeter, $P_i$, to the square root of its surface area $A_i$ [10]:

$$q_j = \frac{P_i}{\sqrt{A_i}}.$$

Within the framework of vertex models of epithelial tissues, this quantity can be interpreted both as a material property of the cells because it is set by the ratio between cell–cell adhesive stress and cell cortical tension, and as an order parameter, which can be used to describe the solid-liquid jamming transition that exists in these tissue driven by cell motility or cell topological rearrangements (cell division, stratification/apoptosis, neighbour exchange, etc.). For a model tissue composed of cells with regular polygonal shapes, this transition has been demonstrated to occur for $q = \langle q_j \rangle = 3.81$, with $\langle \dots \rangle$ denoting an average over all cells in the tissue. Below this threshold, the tissue is in a solid jammed state and above in a fluid-like liquid state. This prediction of the model has been verified in various experimental systems [11], though it is thought that the actual value of q at the transition might be actually higher depending on cell shape anisotropy [12].

##### 3. Second harmonic generation tissue images analysis

Second Harmonic Generation (SHG) signals allow imaging of the collagen contained in the stromal compartment of the esophagus in a label-free fashion. We analysed the spatial organisation of this essential extracellular matrix component in single-plane images taken from z-stacks acquired immediately below the EE basement membrane.

To extract the local orientation of collagen fibres in these images, we followed an approach based on the image structure tensor [13]. This quantity is defined for all pixels in an image as:

$$\boldsymbol{J} = \begin{pmatrix} \langle \partial_x I(x,y),\, \partial_x I(x,y) \rangle_w & \langle \partial_x I(x,y)\,, \partial_y I(x,y) \rangle_w \\ \langle \partial_x I(x,y), \partial_y I(x,y) \rangle_w & \langle \partial_y I(x,y), \partial_y I(x,y) \rangle_w \end{pmatrix}$$

where $I(x,y)$ is the image intensity and $\langle f,g \rangle_w = \iint w(x,y) f(x,y) g(x,y)\, dx\, dy$ is the convolution by a 2D Gaussian kernel $w(x,y)$ of width $\sigma$. Using basic linear algebra, it is possible to determine the eigenvalues and eigenvectors of this tensor, and to define the local dominant orientation θ as the direction of the largest eigenvector of $\boldsymbol{J}$:

$$\theta = \frac{1}{2}\tan^{-1}\left( 2 \frac{\langle \partial_x I(x,y),\, \partial_y I(x,y) \rangle_w}{\langle \partial_y I(x,y),\, \partial_y I(x,y) \rangle_w - \langle \partial_x I(x,y),\, \partial_x I(x,y) \rangle_w} \right)$$

In practice, we used a 2D Gaussian kernel of $\sigma = 6$ pixels width, and the values obtained for θ were locally averaged on a grid with a mesh size of 20 pixels in order to cancel out local variability due to intensity variation and edge effects.

The collagen fibre orientation distributions obtained this way were averaged over at least 3 technical replicates per biological replicate, and data from at least 3 biological replicates per time point were used to plot angular distribution histograms shown in **Extended Data Fig. 5d**. We also characterised for each time point the dominant orientation, the standard deviation of the orientation, and the orientational order parameter of collagen fibre $Q$. This quantity is defined, from the mathematical standpoint, in a similar fashion to the nematic order parameter i.e. as the dominant eigenvalue of the orientation tensor $\boldsymbol{O}$:

$$Q = \sqrt{\langle \cos\theta^{\,2} \rangle + \langle \sin\theta^{\,2} \rangle}.$$

The physical interpretation of this quantity is also similar even if its biophysical origin or the space scales involved are completely different.
